# *COPB2* haploinsufficiency causes a coatopathy with osteoporosis and developmental delay

**DOI:** 10.1101/2020.09.14.297234

**Authors:** Ronit Marom, Lindsay C. Burrage, Aurélie Clément, Bernardo Blanco-Sánchez, Rossella Venditti, Mahim Jain, Ingo Grafe, Daryl A. Scott, Jill A. Rosenfeld, V. Reid Sutton, Marwan Shinawi, Ghayda Mirzaa, Catherine DeVile, Rowenna Roberts, Alistair D Calder, Jeremy Allgrove, Denise G. Lanza, Xiaohui Li, Kyu Sang Joeng, Yi-Chien Lee, I-Wen Song, Joseph M. Sliepka, Dominyka Batkovskyte, Zixue Jin, Brian C. Dawson, Shan Chen, Yuqing Chen, Ming-Ming Jiang, Elda M. Munivez, Alyssa A. Tran, Lisa T. Emrick, David R. Murdock, Neil A. Hanchard, Gladys E. Zapata, Nitesh R. Mehta, Mary Ann Weis, Cole Kuzawa, Abbey Scott, Brenna A. Tremp, Jennifer B. Phillips, Jeremy Wegner, Tashunka Taylor-Miller, Richard A. Gibbs, Donna M. Muzny, Shalini N. Jhangiani, Rolf W. Stottmann, Mary E. Dickinson, John R. Seavitt, Jason D. Heaney, David R. Eyre, Catherine G. Ambrose, Undiagnosed Diseases Network Monte Westerfield, Maria Antonella De Matteis, Brendan Lee

**Author notes:** MJ: Kennedy Krieger Institute, Baltimore MD, USA. KSJ: Department of Orthopaedic Surgery, University of Pennsylvania, Philadelphia PA, USA. IG: Division of Endocrinology, Diabetes, and Bone Diseases, Department of Medicine III and Center for Healthy Aging, University Clinic Dresden, Germany; Center for Regenerative Therapies Dresden, University Dresden, Germany. DB: Department of Molecular Medicine and Surgery, Karolinska Institutet, Stockholm, Sweden. To whom correspondence should be addressed: Brendan Lee, M.D., Ph.D., Department of Molecular and Human Genetics, Baylor College of Medicine, One Baylor Plaza, Room R814, Houston, TX 77030.

## Abstract

Coatomer complexes function in the sorting and trafficking of proteins between subcellular organelles. Pathogenic variants in coatomer subunits or associated factors have been reported in multi-systemic disorders, i.e., coatopathies, that can affect the skeletal and central nervous systems. We have identified loss-of-function variants in *COPB2*, a component of the coatomer complex I (COPI), in individuals presenting with osteoporosis, fractures and developmental delay of variable severity. Because the role of COPB2 in bone has not been characterized, we studied the effect of *COPB2* deficiency on skeletal development in mice and zebrafish. *Copb2*^*+/−*^ mice showed low bone mass and decreased bone strength. In zebrafish, larvae carrying a *copb2* heterozygous frameshift variant showed delayed mineralization. *copb2*-null embryos showed endoplasmic reticulum (ER) and Golgi disorganization, and embryonic lethality. *COPB2* siRNA-treated fibroblasts showed delayed collagen trafficking with retention of type I collagen in the ER and Golgi, and altered distribution of Golgi markers. Our data suggest that *COPB2* haploinsufficiency leads to disruption of intracellular collagen trafficking and osteoporosis, which may improve with ascorbic acid supplementation. This work highlights the role of COPI complex as a critical regulator of bone mass and identifies a new form of coatopathy due to *COPB2* deficiency.

## Introduction

Vesicle coat proteins are an essential and evolutionarily conserved group of proteins that form the molecular machinery responsible for the sorting and trafficking of proteins and lipids within the cell (1–3). Pathogenic variants in genes encoding subunits of coat complexes (coatomers), or in accessory and regulatory factors important for their function, have been implicated in a number of genetic disorders collectively termed coatopathies (2). Disruption of secretory pathways can cause retention of cargo proteins within the endoplasmic reticulum (ER) and Golgi compartments, ultimately leading to impaired biogenesis and function of these organelles, ER stress, and decreased cell viability. Abnormal vesicular transport also affects post-translational modifications (specifically glycosylation) of secreted proteins at the Golgi complex and may alter the extracellular matrix composition. Although this can affect multiple organ systems, the central nervous system is most frequently affected, with individuals typically manifesting with microcephaly and developmental delay (2, 4, 5).

Skeletal development and bone strength are highly dependent on the proper biosynthesis, post-translational modifications, assembly, and cross-linking of the collagen fibrils (6), including type I collagen within the bone and type II collagen within the cartilage. Procollagen (PC) molecules undergo multiple post-translational modifications while being transported through the ER, Golgi complex, and the plasma membrane (7). The ER export of fibrillar PC, such as type I and type II procollagen (PCI and PCII), requires the coordinated action of PC receptors such as transport and Golgi organization (TANGO) /cutaneous T cell lymphoma-associated antigen 5 (cTAGE5) proteins, coat protein complex II (COPII) and transport protein particle (TRAPP) complex components, and the retrograde recruitment of coat protein complex I (COPI)-coated ERGIC53-containing vesicles (3, 8–10). Abnormal export of PCI and PCII from the ER to the Golgi and plasma membrane due to defects in vesicular trafficking components has been associated with skeletal dysplasias (11–15).

The COPI complex plays a critical role in protein transport within the Golgi complex and between the ER and Golgi (16–18). COPI subunits are essential for notochord development in zebrafish (19). In humans, geroderma osteodysplasticum is caused by loss of the COPI scaffold protein GORAB (20), and defects in *ARCN1*, encoding COPI complex subunit, are associated with syndromic rhizomelic short stature (21).

Here we report heterozygous, loss-of-function variants in *COPB2*, a component of the COPI coatomer complex, in six individuals from five unrelated families presenting with a clinical spectrum of osteoporosis or osteopenia, with or without fractures, and developmental delay of variable severity. A hypomorphic, homozygous missense variant in *COPB2* was previously reported in two siblings with microcephaly, spasticity and developmental delay (22), in whom we also identified low bone mass. We focused on the effect of *COPB2* deficiency on bone development and demonstrate that pathogenic variants in *COPB2* lead to early onset osteoporosis, and that COPB2 and the COPI complex are essential regulators of bone mass.

## Results

### Clinical presentation and molecular studies

We have identified 6 individuals (Subjects 1-6) with overlapping phenotypes who carry putatively deleterious variants in *COPB2.* Their molecular and clinical finding are summarized below and in Table 1.

**Table 1:** Clinical summary and *COPB2* variants identified.

|  | Subject 1 (Family 1) | Subject 2 (Family 2) | Subject 3 (Family 3) | Subject 4 (Family 4) | Subject 5 (Family 5) | Subject 6 (Family 5 ) |
| --- | --- | --- | --- | --- | --- | --- |
| Gender | F | F | M | F | M | F |
| Age at most recent evaluation (years) | 9 | 9 | 4 | 3 | 9 | 12 |
| Ethnicity | Caucasian/Native American | Caucasian/Native American | Caucasian | Asian | Caucasian/Native American | Caucasian/Native American |
| *COPB2* Variant (hg19, NM_004766.2) | c.1237_1238delAA<br>p.Lys413AspfsTer3<br>g.139088354_355delTT<br>(exon 11/22) | c.1906dupA<br>p.The636AsnfsTer7<br>g.139081338dupT<br>(exon 16/22) | c.1206-2A>G<br>g.139088388T>C<br>(IVS 10) | c.247dupG<br>p.Val83GlyfsTer14<br>g.139097997dupC<br>(exon 4/22) | c.760C>T p.Arg254Cys<br>g.139092642G>A<br>(exon 8/22) | c.760C>T p.Arg254Cys<br>g.139092642G>A<br>(exon 8/22) |
| Inheritance | *De novo* (heterozygous) | Not maternal (heterozygous) | *De novo* (heterozygous) | *De novo* (heterozygous) | Homozygous | Homozygous |
| **General** |  |  |  |  |  |  |
| Prematurity | No (39 weeks) | Yes (33 weeks) | No (stated as term) | No (40 weeks) | Yes (36 weeks) | Yes (35 weeks) |
| IUGR or small for age at birth | No | No | No | No | No | No |
| Growth parameters on most recent evaluation (percentile) | Weight 27 %ile<br>Height 35 %ile | Weight 43 %ile<br>Height 10 %ile | Weight 5 %ile<br>Height 1%ile | Weight 25 %ile<br>Height 70 %ile | Weight 63 %ile<br>Height 1 %ile | Weight 1 %ile<br>Height 30 %ile |
| Head circumference (Z score) | 0.65 | -2.19 | -0.31 | -4 | -8 | -5.5 |
| **Musculoskeletal Features** |  |  |  |  |  |  |
| Osteopenia | Yes | Yes | Yes | Unknown | Yes | Yes |
| Fractures | Yes | No | Yes | No | No | No |
| **Neurological Features** |  |  |  |  |  |  |
| Developmental delay | Yes | Yes | Yes | Yes | Yes | Yes |
| Intellectual disability | No, but has learning disability & ADHD | Yes, mild | N/A | N/A | Yes, severe (nonverbal) | Yes, severe (nonverbal) |
| Spasticity (ambulatory status) | No (normal gait) | Yes (wide-based unsteady gait, progressed to require a walker and wheelchair) | No (wide-based gait) | Possible in ankles bilaterally (mild, with normal gait) | Yes (non ambulatory) | Yes (non ambulatory) |
| Seizures | No | No | No | No | Yes | Yes |
| Brain MRI | Focal cortical dysplasia | Normal | Normal | Normal | microcephaly, simplified gyral pattern, thin corpus callosum, delayed myelination | microcephaly, simplified gyral pattern, thin corpus callosum, delayed myelination |
| **Prior Genetic Testing** |  |  |  |  |  |  |
| Copy number variant analysis | Negative | Copy number gain on chromosome 12p12.1 (24,692,382-25,353,987; hg19), maternally inherited | Negative | Negative | A 16.8 Mb region of absence of heterozygosity (AOH) on chromosome 3q22.1-3q24 (131374802-148170986; hg19) | A 16.8 Mb region of absence of heterozygosity (AOH) on chromosome 3q22.1-3q24 (131374802-148170986; hg19) |
| Other clinical testing | *COL1A1/COL1A2* sequencing and recessive OI gene panel – negative, metabolic labs – normal, CDG – negative, clinical ES – negative | Hereditary spastic paraplegia panel- negative, metabolic labs and CSF studies – normal, CDG – negative, clinical ES – negative | OI dominant gene panel (*COL1A1/COL1A2/IFITM5*) - negative | Autism/ID gene panel - negative |  | Primary microcephaly gene panel- negative |

Abbreviations: CDG – congenital disorders of glycosylation, CSF- cerebrospinal fluid, ES-exome sequencing, ID- intellectual disability, N/A- not applicable, OI- osteogenesis imperfecta.

***Subject 1*** is a 9-year-old female who presented with her first fracture at 2 years of age, followed by more than 10 fractures of long bones. Bone densitometry via dual energy x-ray absorptiometry (DXA) scan at 3 years of age indicated lumbar spine Z score of −3.2. Bisphosphonate therapy was initiated, and a repeat DXA scan at 5 years of age showed normal bone mineral density (BMD; lumbar spine Z score of 0.2; left hip Z score of 1.8; right hip Z score of 2.8). Over time, the frequency of her fractures decreased, but she continued to suffer chronic bone pain. Her growth was normal (height at the 60th percentile). She did not have dentinogenesis imperfecta, blue sclera, joint hypermobility, bone deformities or scoliosis. There was no family history of osteopenia or recurrent fractures. Her development was significant for speech delay, attention deficit hyperactivity disorder (ADHD) and learning disabilities. Brain MRI showed focal cortical dysplasia of left frontal gyrus although she had no seizures. Previous genetic evaluation included normal *COL1A1/COL1A2, CRTAP, LEPRE1, FKBP10, PPIB, SERPINF1, SERPINH1*, and *SPF7* targeted sequencing. SNP-based copy number variant (CNV) analysis did not detect pathogenic deletions or duplications. Metabolic evaluations, including plasma amino acids, urine amino acids, an acylcarnitine profile, a lipid panel and a congenital disorders of glycosylation panel, were normal. Research trio exome sequencing did not detect variants in genes known to cause osteogenesis imperfecta (OI) or other low-bone mass phenotypes. Further analysis of the data identified a heterozygous c.1237_1238delAA, p.Lys413Aspfs*3 (NM_004766.2) variant in *COPB2*. This variant has not been reported in gnomAD (http://gnomad.broadinstitute.org) (23) and is predicted to cause loss of function due to a shift in the reading frame. Sanger sequencing of DNA from the proband and her parents confirmed that the variant is *de novo*. mRNA sequencing from the proband’s whole blood demonstrated that this variant leads to nonsense-mediated mRNA decay (**Supplemental figure 1**). Real-time qPCR analysis in lymphoblastoid cells derived from Subject 1 supported this finding by showing significantly reduced expression of *COPB2*, as compared to her parents and sibling (**Figure 1A**).

**Figure 1.**
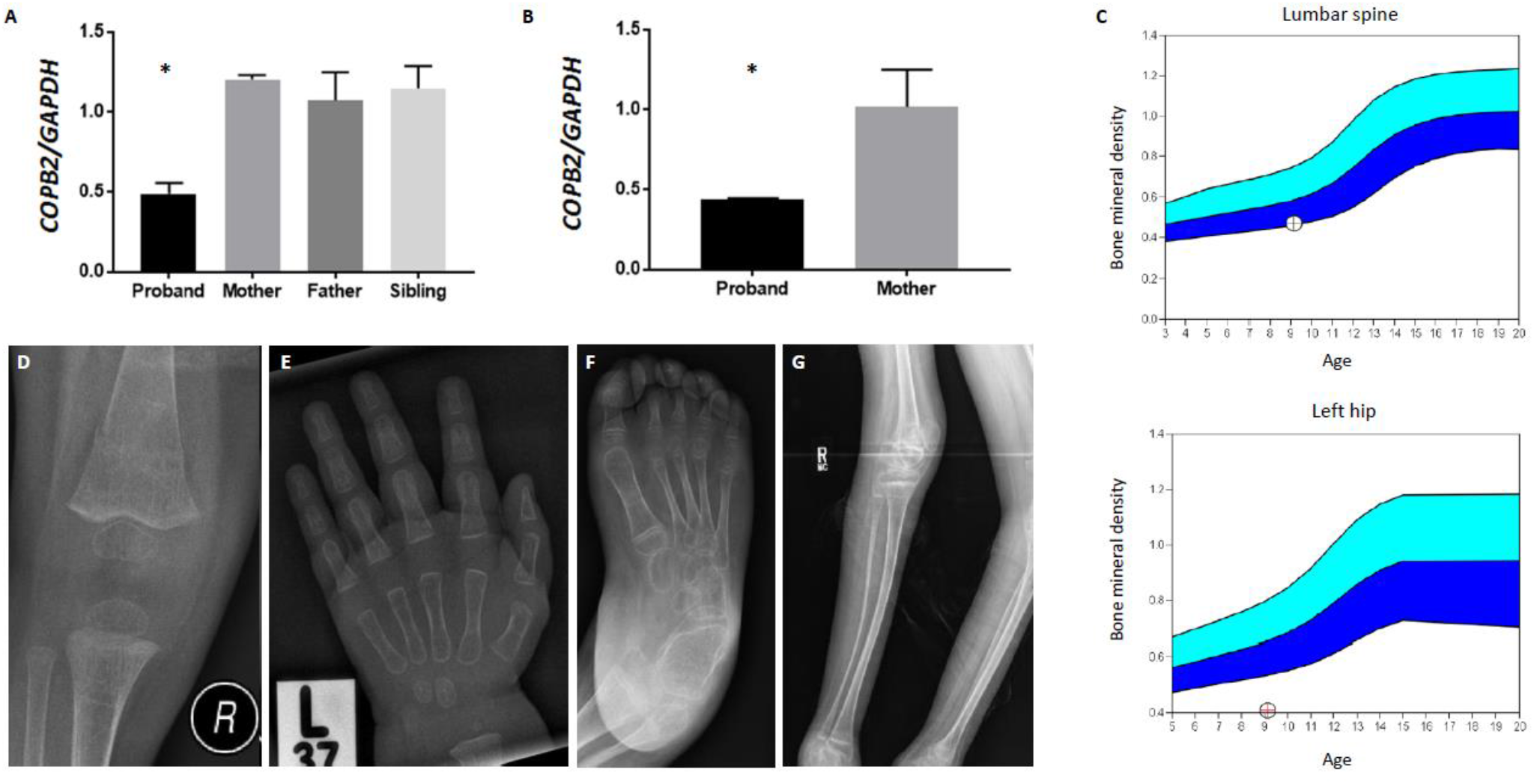
*COPB2* variants lead to osteopenia and fractures. **(A)** *COPB2* qPCR in lymphoblastoid cells from Subject 1 (proband) showing decreased *COPB2* expression at around 50% compared to expression in parents and sibling. **(B)** *COPB2* qPCR in skin fibroblasts from Subject 2 (proband) and her mother showing decreased *COPB2* expression at around 50% compared to expression in parent. Results shown as *COPB2/GAPDH* relative mRNA expression, p= 0.0002 (one-way ANOVA) for subject 1, and p=0.002 (t test) for subject 2. **(C)** Bone densitometry scan in Subject 2, showing low bone mineral density of the lumbar spine (upper panel) and left hip (lower panel). **(D)** Right knee radiograph of Subject 3 at age 14 months showing fracture of distal femoral metadiaphysis and osteopenia; **(E)** Left hand radiograph of Subject 3 at age 20 months showing thin metacarpal cortices; **(F)** Right foot radiograph of Subject 5 at age 8 years showing diffuse osteopenia and hind-foot varus deformity; **(G)** Right leg radiographs of Subject 6 at age 12 years showing gracile, over-tubulated long bones;

***Subject 2*** is a 9-year-old female that presented with global developmental delay, microcephaly and progressive spastic paraparesis. She was initially ambulatory but had wide-based gait and started developing spasticity at 2-3 years of age. She is able to walk a few steps by herself but requires a walker or wheelchair to travel longer distances. She does not have a history of fractures, but a DXA scan performed at 9 years of age showed a total spine Z score of −1.9 and femoral neck Z score of −2.4 (**Figure 1C**). The family history is significant for a sibling who has severe developmental delay and is non-ambulatory due to a large, *de novo* 8q21.3-q24.3 duplication that is 52.42 Mb in size. There is no family history of osteopenia or recurrent fractures. MRI of brain and spine at 2 years of age showed scoliosis of the thoracolumbar spine but was otherwise normal. Electromyography and nerve conduction velocity studies were normal. Previous genetic evaluation included a normal hereditary spastic paraplegia gene panel. Array-based CNV analysis revealed a maternally inherited 12p12.1 duplication that is 662 kb in size (including *SOX5*, *BCAT1* and *LRMP* genes) that was considered an unlikely explanation for her phenotype. Clinical exome sequencing revealed a heterozygous c.1906dupA, p.Thr636Aspfs*7 (NM_004766.2) variant in *COPB2* that was classified as a variant of uncertain significance. Subsequently, the family consented for transfer of those data to study in the Undiagnosed Diseases Network (UDN). The variant in *COPB2* is novel and not reported in gnomAD (23). It is predicted to cause loss of function due to a shift in the reading frame. The mother and full siblings tested negative for this variant, but her deceased father was not available for testing. Real-time qPCR analysis in fibroblast cells derived from Subject 2 showed significantly reduced expression of *COPB2*, as compared to her mother (**Figure 1B**).

***Subject 3*** is a 4-year-old male who first presented with fracture of the right distal femoral metadiaphysis at 14 months of age (**Figure 1D**). He had another fracture of the right distal femur at 20 months due to minor trauma. A skeletal survey demonstrated recurrent insufficiency fractures of the distal right femur, generalized osteopenia (**Figure 1D, E**), varus bowing deformities of both ulnae, no rachitic changes, apparent cavo-varus deformities of the hind feet, and Wormian bones at the lambdoid suture (the latter could be a normal variant). His growth was notable for short stature (height Z score at – 2.3). His exam was significant for pale grey sclerae, normal dentition, low muscle tone and mild joint hypermobility. A kyphotic posture was noted while sitting although the spine was straight. There was no family history of osteopenia or recurrent fractures. He is the seventh of eight children born to non-consanguineous Ashkenazi Jewish parents. His parents and siblings are all healthy. His development was significant for gross motor delay. A brain MRI was normal. An array-based CNV analysis and a dominant osteogenesis imperfecta gene panel, including *COL1A1*, *COL1A2* and *IFITM5* were negative. The patient was recruited into the 100,000 genomes project (UK Department of Health funded genomes project) and genome sequencing identified a *de novo* variant, c.1206-2A>G (NM_004766.2), in *COPB2* that is novel and not reported in gnomAD (23). The variant is predicted to affect splicing.

***Subject 4*** is a 3-year-old female who presented with global developmental delay, abnormal muscle tone, poor weight gain and microcephaly. Prenatal history was unremarkable, and fetal growth measurements were within the normal limits. Head circumference growth began declining at 4 months of life from 2 standard deviations (SD) below the mean to 2.86 SD below the mean at age 1 year 4 months, and to 4 SD below the mean at 3 years of age. Her medical history was also remarkable for atopic dermatitis, mild joint hypermobility, and feeding issues. She does not have a history of fractures and has not had skeletal radiographs to evaluate for osteopenia. A brain MRI performed at 2 years of age was normal. A SNP-based CNV analysis was negative. An Autism/Intellectual Disability XPanded Panel (GeneDx) that revealed a *de novo*, heterozygous variant, c.247dupG, p.V83Gfs*14, in *COPB2* (NM_004766.2) that is novel and not reported in gnomAD (23). This frameshift variant is predicted to result in protein truncation or nonsense-mediated mRNA decay.

***Subject 5***, a 9-year-old male and ***Subject 6***, a 12-year-old female are siblings that were described in a prior publication (22). Their bone phenotype had not been evaluated at the time of the previous report. They presented with global developmental delay, spasticity, seizures and severe microcephaly. They do not have a history of fractures, but skeletal radiographs showed gracile, over-tubulated bones and osteopenia (**Figure 1F, G**). Brain MRI findings in both siblings demonstrated microcephaly with simplified gyral pattern, thin corpus callosum and delayed myelination. A primary microcephaly gene panel in Subject 6, including *ASPM*, *CDK5RAP2*, *CENPJ*, *MCPH1*, and *STIL*, was negative. Array-based CNV analysis identified a 16.8 Mb region of absence of heterozygosity on chromosome 3q22.1-q24 (chr3: 131,374,802-148,170,986; hg19). A subsequent research exome sequencing analysis indicated a shared recessive homozygous missense variant, c.760C>T, p.Arg254Cys (NM_004766.2) in *COPB2*. This variant was reported 3 times in gnomAD (23) in European (non-Finnish) population (allele frequency 0.00001063) but only in a heterozygous status.

### Copb2^+/−^ mice exhibit low bone mass and decreased bone strength

To better understand the clinical relevance of these findings, a mouse carrying a *Copb2* null allele was generated using CRISPR/Cas9 technology (**Supplemental Figure 2**). In crosses of *Copb2*^+/−^ mice, no *Copb2*^*−/−*^ pups were identified postnatally. Homozygosity for a similar CRISPR/Cas9-generated allele (*Copb2*^*em1(IMPC)Bay*^) produced by the Baylor College of Medicine (BCM) component of the Knockout Mouse Production and Phenotyping (KOMP2) project was found to be embryonic lethal by E18.5 (http://www.mousephenotype.org/). Therefore, phenotypic analysis was performed using *Copb2*^*+/−*^ mice. Skeletal radiographs of hindlimbs at 2 months of age did not reveal gross bone deformities. Analysis of bone mass using micro-CT (μCT) imaging showed a 15-20% reduction in spine bone volume/total volume (BV/TV) in 2-month-old *Copb*^*+/−*^ male and female mice compared to control wild-type littermates (**Figure 2A**). In females, the spinal trabecular number, trabecular thickness, and femur cortical thickness were significantly reduced as compared to wild-type littermates (**Figure 2A, B**).

**Figure 2.**
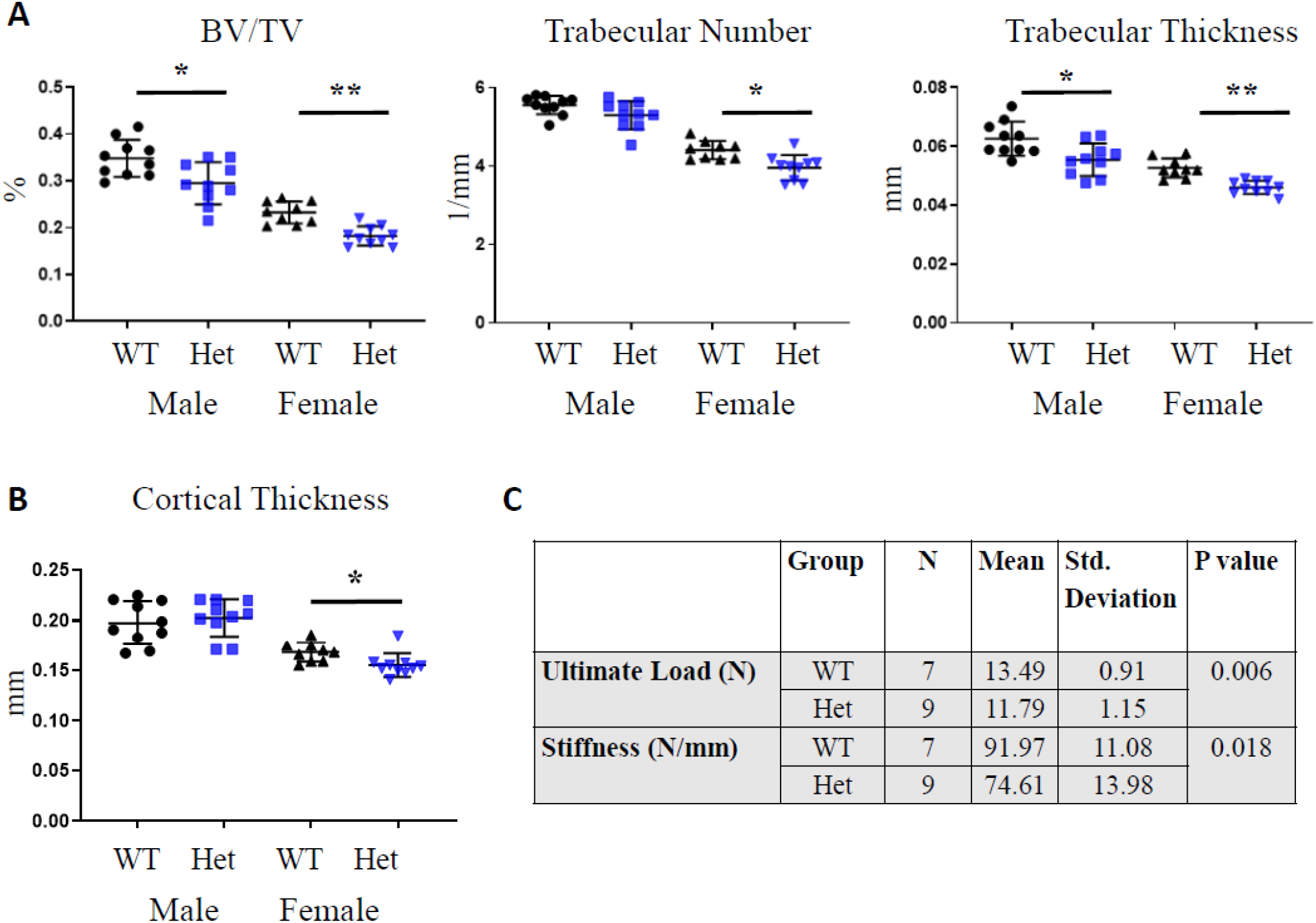
*Copb2* heterozygous mice exhibit low bone mass phenotype. **(A)** Micro-CT analysis of spine showed significant decrease in bone architectural parameters in *Copb2*^*+/−*^ mice (Het), including the bone volume/total volume (BV/TV, p=0.014 for males, p=0.0006 for females) and trabecular thickness (p=0.007 for males, p<0.0001 for females). Females also showed decreased trabecular number (p= 0.002). **(B)** Micro-CT analysis of femurs showed mild reduction in cortical thickness in the females (p=0.003). Mean values ± SD are indicated for both (A) and (B) (* p<0.05, ** p< 0.001). **(C)** Biomechanical testing of femurs by 3-point bending demonstrated decreased bone strength, with reduced maximal load and stiffness in femurs of *Copb2*^+/−^ female mice (Het), consistent with osteopenia.

Three-point bending was performed in *Copb2*^*+/−*^ mice femurs to assess bone strength. *Copb2*^*+/−*^ bones showed decreased ultimate load and stiffness, indicating reduced bone strength (**Figure 2C**). However, the ultimate strength and elastic modulus were comparable to wild-type littermates (**Supplemental Table 1**). Notably, there was no increase in bone brittleness in femurs of the *Copb2*^*+/−*^ mice (no significant difference in plastic displacement), indicating that COPB2 deficiency does not lead to abnormal bone material mechanical properties, as would be observed in OI caused by structural variants or altered post-translational modifications of type I collagen (24).

To understand the cellular consequences of COPB2 deficiency in bone, histomorphometric analysis in L4 vertebrae of 2-month-old mice was performed. However, this did not reveal significant differences in osteoblast and osteoclast numbers, or in the bone formation rate or mineralization rate between *Copb2*^*+/−*^ mice and wild-type littermates (**Supplemental Figure 3**). *In vitro* differentiation of mouse calvarial cells, as measured by alizarin red staining for mineralization and alkaline phosphatase activity, did not significantly differ between *Copb2*^*+/−*^ mice and their wild-type littermates (**Supplemental Figure 3**). *Col1a1* expression, as measured by qPCR in RNA derived from bone tissue and cell cultures did not significantly differ between *Copb2*^*+/−*^ and wild-type littermates (**Supplemental Figure 4**). Additionally, mouse femurs were analyzed for possible collagen over-modification to check for abnormal processing of type I collagen, which is characteristic of OI caused by qualitative defects in type I collagen, or by pathogenic variants in components of the collagen 3-hydroxylation complex (25). This analysis did not reveal significant changes in lysine or proline modification (**Supplemental Figure 4**).

### copb2 mutant zebrafish embryos show delayed mineralization, abnormal secretion of type II collagen, and disruption of the Golgi and ER

To study the developmental consequences of *COPB2* loss of function in a second animal model, a mutant allele of the zebrafish ortholog, *copb2*, was generated using CRISPR/Cas9 technology by introducing a frameshift followed by a premature stop codon (*copb2*^*b1327*^, p.K10Tfs*11; **Supplemental Figure 5**). Consistent with a previously analyzed zebrafish null allele of *copb2* (19, 26), *copb2*^*b1327/b1327*^ homozygous mutant animals showed a severe phenotype at 24 hours post fertilization (hpf), including: kinked notochord, pigmentation defect, hydrocephaly, thinner midbrain-hindbrain boundary and embryonic lethality. In the developing skeleton, alizarin red-stained wild-type larvae exhibit calcified bones at 7 days post fertilization (dpf), including the operculum (**Figure 3A**). A fraction of *copb2*^*b1327/+*^ larvae showed absence of calcification of the opercula (4/71 larvae; **Figure 3B, C**). There was no difference in the intensity of alizarin red staining between wild-type and *copb2*^*b1327/+*^ heterozygous mutant juveniles at 34 dpf (**Figure 3D**), suggesting that the defect in operculum calcification is due to delayed mineralization. No differences in Alcian blue staining for cartilage were detected (**Figure 3A, B**).

**Figure 3.**
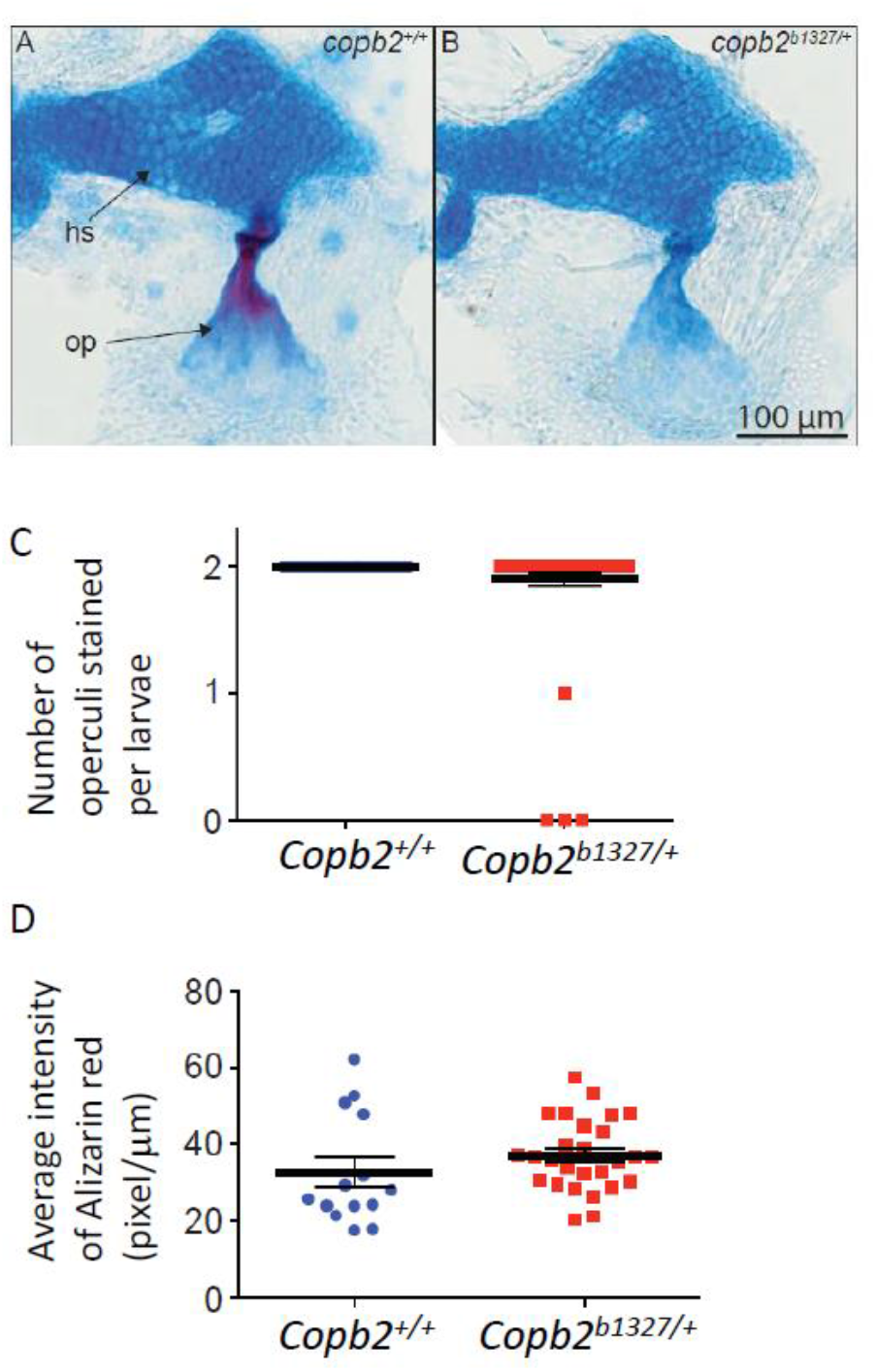
Calcification of the operculum is delayed in heterozygous *copb2b*^*1327/+*^ mutant zebrafish. Alcian blue/alizarin red staining of 7 dpf *copb2* wild-type **(A)** and *copb2*^*b1327/+*^ heterozygous **(B)** mutant larvae (hs: hyosymplectic; op: operculum). **(C)** Number of calcified opercula in 7 dpf *copb2* wild-type (n=34 larvae) and *copb2*^*b1327/+*^ heterozygous mutant larvae (n=71 larvae). **(D)** Average intensity of alizarin red per operculum of 34 dpf *copb2* wild-type (n=14 juveniles) and *copb2*^*b1327/+*^ heterozygous mutants (n=27 juveniles). Data are presented as mean ± SEM.

In *copb2*^*b1327/b1327*^ homozygous mutant embryos, the ER and the Golgi apparatus appeared mislocalized and fragmented (**Figure 4A-D**) as observed previously (19). Defects in type II collagen secretion with cytoplasmic aggregation in notochord cells were observed (**Figure 4E, F**) supporting the evolutionarily conserved role of the Golgi complex in collagen trafficking. Although *copb2*^*b1327/b1327*^ zebrafish exhibit hydrocephaly and a thinner midbrain-hindbrain boundary, sections through the brain did not reveal any other obvious abnormalities (**Supplemental Figure 6**). Likewise, *Copb2*^−/−^ mouse embryos did not show microcephaly or brain anatomical abnormalities, and the heterozygous mice had no observed neurological phenotype (http://www.mousephenotype.org/). Interestingly, developmental delay, with or without microcephaly, had been reported in coatopathies and other disorders related to vesicular transport dysfunction (2, 4, 5).

**Figure 4.**
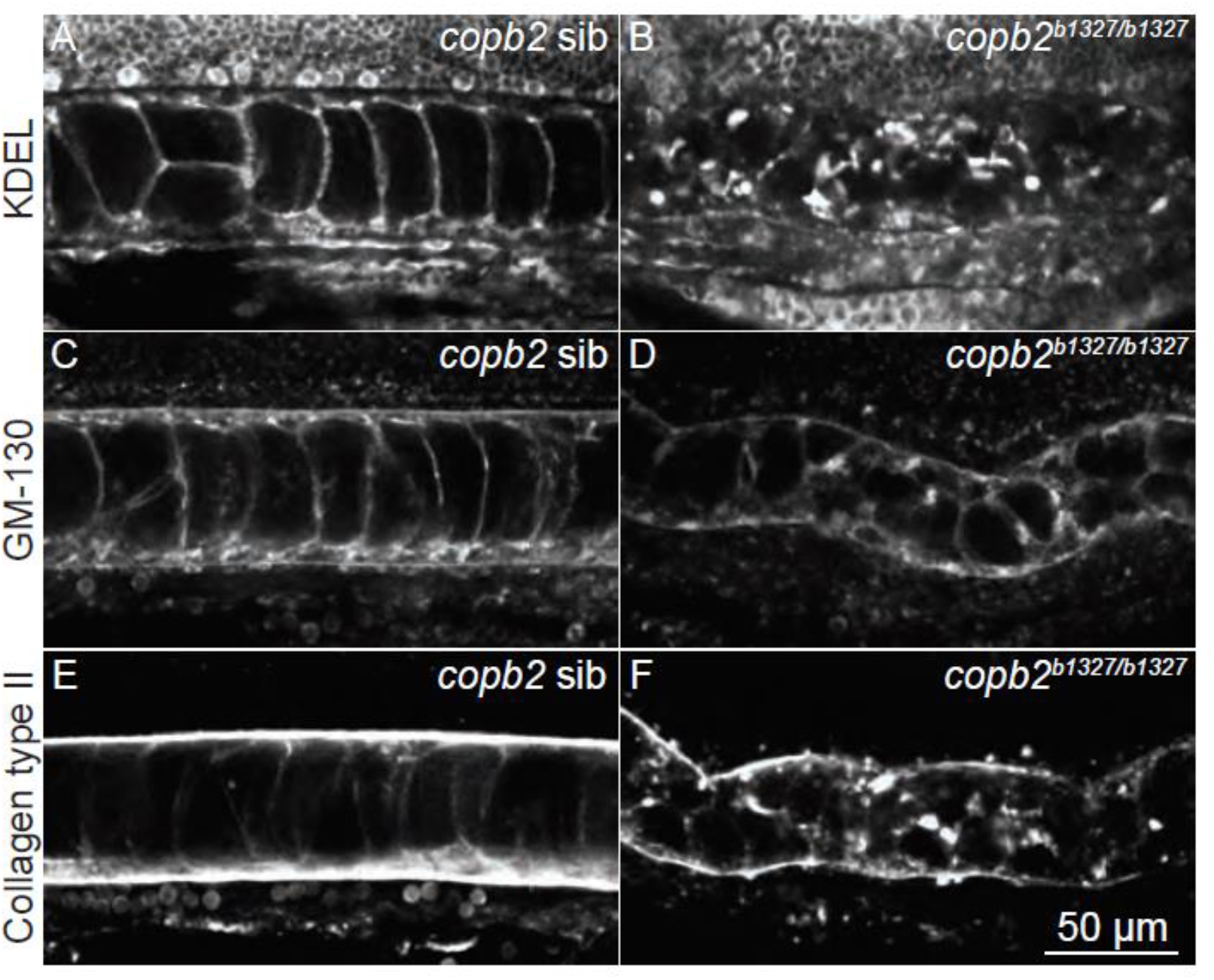
Structural integrity of the ER and Golgi apparatus and protein secretion are compromised in *copb2*^*b1327/b1327*^ zebrafish mutants. **(A-F)** Confocal sections of the notochord. Immunolabeling of KDEL in *copb2* siblings **(A)** and *copb2*^*b1327/b1327*^ mutant **(B)** embryos. Immunolabeling of GM130 in *copb2* siblings **(C)** and *copb2*^*b1327/b1327*^ mutant **(D)** embryos. Immunolabeling of type II Collagen in *copb2* siblings **(E)** and *copb2*^*b1327/b1327*^ mutant **(F)** embryos (anterior is to the left and dorsal to the top). Images were taken at the level of the yolk extension at 30 hpf.

### COPB2 depletion is associated with delayed collagen trafficking and disorganization of the Golgi complex

To better understand the molecular consequences of COPB2 deficiency, the organization of the ER-Golgi intermediate compartment (ERGIC) and Golgi complex, and the trafficking of different classes of cargo including PCI were assessed by immunofluorescence microscopy in human fibroblasts or Hela cells, where the expression of *COPB2* was titrated by siRNA treatment (**Figure 5 and Supplemental Figure 7**). *COPB2* siRNA-treated cells exhibited a disorganized distribution of Golgi markers that was proportional to the extent of the COPB2 depletion (**Figure 5A and Supplemental Figure 7A, B**). Additionally, a marked reduction of the punctate peripheral pattern of the ERGIC, as assessed by COPI, GBF1 and KDEL receptor labeling, was induced by COPB2 depletion suggesting impaired Golgi-to-ER retrograde trafficking (**Figure 5B and Supplemental Figure 7C**).

**Figure 5.**
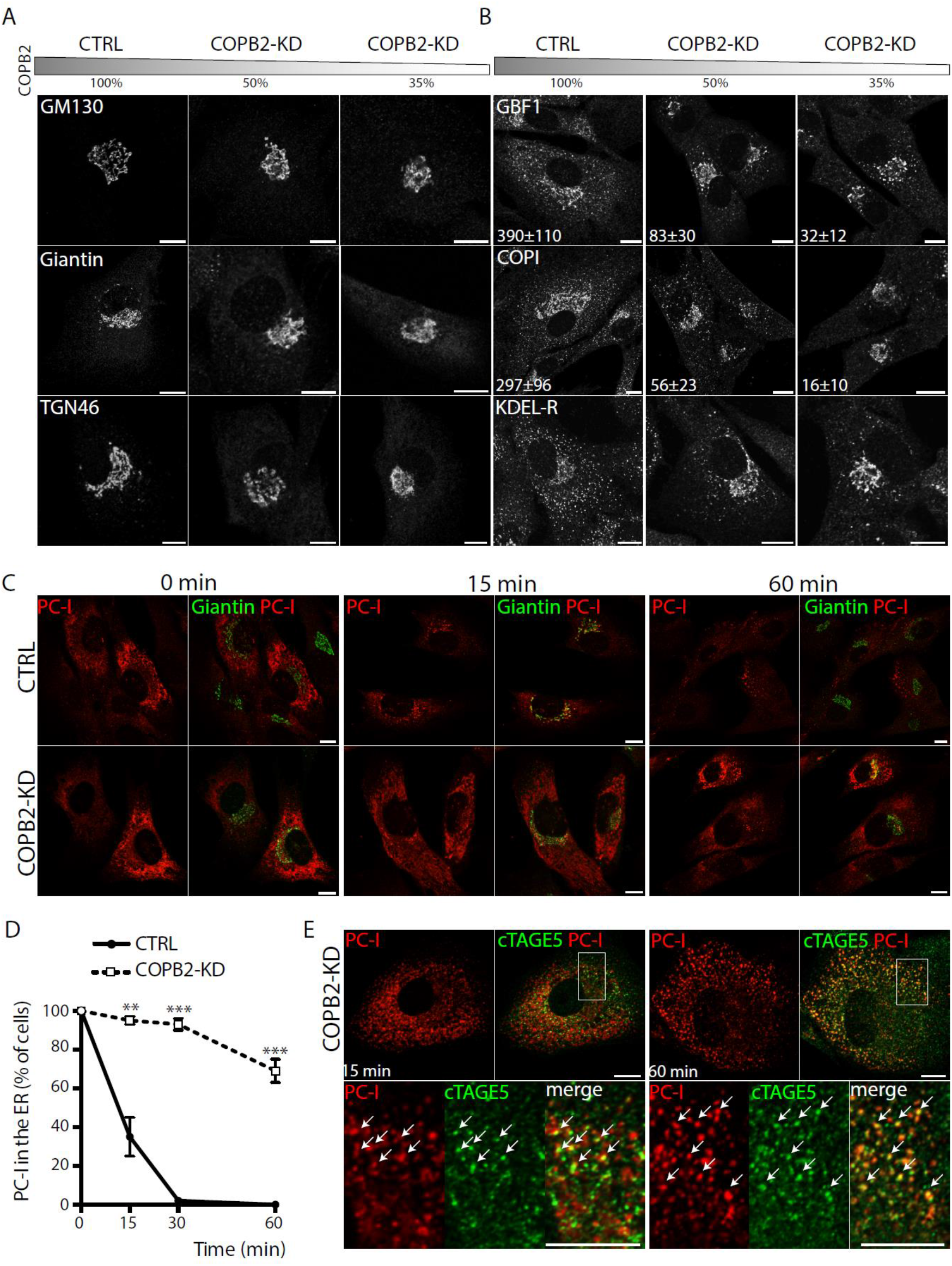
Partial depletion of COPB2 induces disorganization of the Golgi complex and impairs ER exit of PCI in human fibroblasts. (**A**) BJ-5ta human fibroblasts (HFs) were mock (CTRL) or COPB2-siRNA treated for 36 hours to obtain different levels of COPB2 reduction (50% and 35% using 20 nM and 100 nM siRNAs, respectively). The gray bar indicates the residual COPB2 protein level after siRNA treatment. Cells were immunolabeled using an antibody against the cis-Golgi marker GM130 (upper panel), medial Golgi marker Giantin (middle panel) and trans-Golgi protein TGN46 (lower panel). Scale bars=10 μm. (**B**) Immunofluorescence analyses of CTRL and COPB2-KD HFs for several proteins of the ER-Golgi intermediate compartment (ERGIC): GBF1, COPI (evaluated using an anti-coatomer mAbCM1A10) and KDEL receptor (all markers in gray). Scale bars, 10 μm. Numbers represent quantification of GBF1-positive and COPI-positive spots per cell from one representative experiment. N=3 experiments, n=50 cells counted, ± S.D. (**C**) PCI transport in human fibroblasts. CTRL and COPB2-KD (50% of residual protein level) cells were shifted to 40°C for 3 hours to accumulate PCI (red) in the ER. Cells were then either fixed immediately (0 min) or incubated for different times (15 and 60 minutes) at 32°C in the presence of ascorbate and CHX. During the temperature block, both CTRL and COPB2-KD cells accumulate PCI into the ER. In control cells, PCI co-localizes with the Golgi marker Giantin (green) after 15 minutes, but does not in COPB2-KD cells where it fails to exit the ER. Scale bars=10 μm. (**D**) Quantification of the ER exit of PCI in CTRL and COPB2-KD HFs expressed as percentage of cells with PCI in the ER. N=3 experiments, n=100 cells counted. Mean values, ± SD (**p<0.001, *** p<0.0001). (**E**) PC-I transport assay in COPB2-KD human fibroblasts as in C. Cells were immunolabeled for PCI and cTAGE5 as a marker of the ER exit sites. PCI co-localized with cTAGE5 15 and 60 minutes after the release. Bottom panels are enlargements of the boxed areas with white arrows indicating the co-localizing PCI with cTAGE5. Scale bars=10 μm.

To analyze the impact of COPB2 depletion on intracellular trafficking of newly synthesized cargos, we used a temperature-based synchronization protocol that depends on the temperature sensitivity of PCI folding that is retained in the ER at 40°C (12). We found that *COPB2* siRNA-treated fibroblasts had a marked delay in ER export of PCI, which was retained in the ER (**Figure 5C, D**) at the level of ER exit sites (as labelled by cTAGE5, **Figure 5E**) for up to 60 minutes after the release of the 40°C temperature block. By contrast, transport of a temperature-sensitive variant of vesicular stomatitis virus G protein (VSV-G) that served as a control cargo was only slightly delayed (**Supplemental Figure 7**), suggesting a preferential requirement for COPB2 in the intracellular transport of PCI.

### Ascorbic acid rescues the phenotype of copb2-deficient zebrafish

Ascorbic acid (vitamin C) serves as a cofactor for PC hydroxylases and induces the intracellular trafficking and secretion of collagen molecules (7, 16, 27). Given the essential role of ascorbic acid in PC metabolism, the effect of ascorbic acid supplementation was studied in *COPB2*-deficient animal models. Ascorbic acid treatment rescued type II collagen secretion in *copb2*^*b1327/b1327*^ homozygous zebrafish mutant embryos in a dose-dependent manner (**Figure 6B, D, F, G**). A partial rescue of collagen type II secretion was observed in some *copb2*^*b1327/b1327*^ homozygous mutant embryos treated with 100 mM ascorbic acid (29% of mutants; **Figure 6B,D,G**). A full rescue (33% of mutants) was observed when *copb2*^*b1327/b1327*^ homozygous mutant embryos were treated with 200 mM Ascorbic acid (**Figure 6F,G**). Additionally, the morphological defect of the notochord was rescued in embryos in which secretion of type II collagen was improved (**Supplemental Figure 8B, D, F**). A trend of improvement in BMD was detected by μCT analysis of BV/TV in *Copb2*^*+/−*^ mice that received ascorbic acid-enriched diet; however, this effect may have been blunted by the fact that mice, unlike humans and zebrafish, can endogenously synthesize ascorbic acid (**Supplemental Figure 9**).

**Figure 6.**
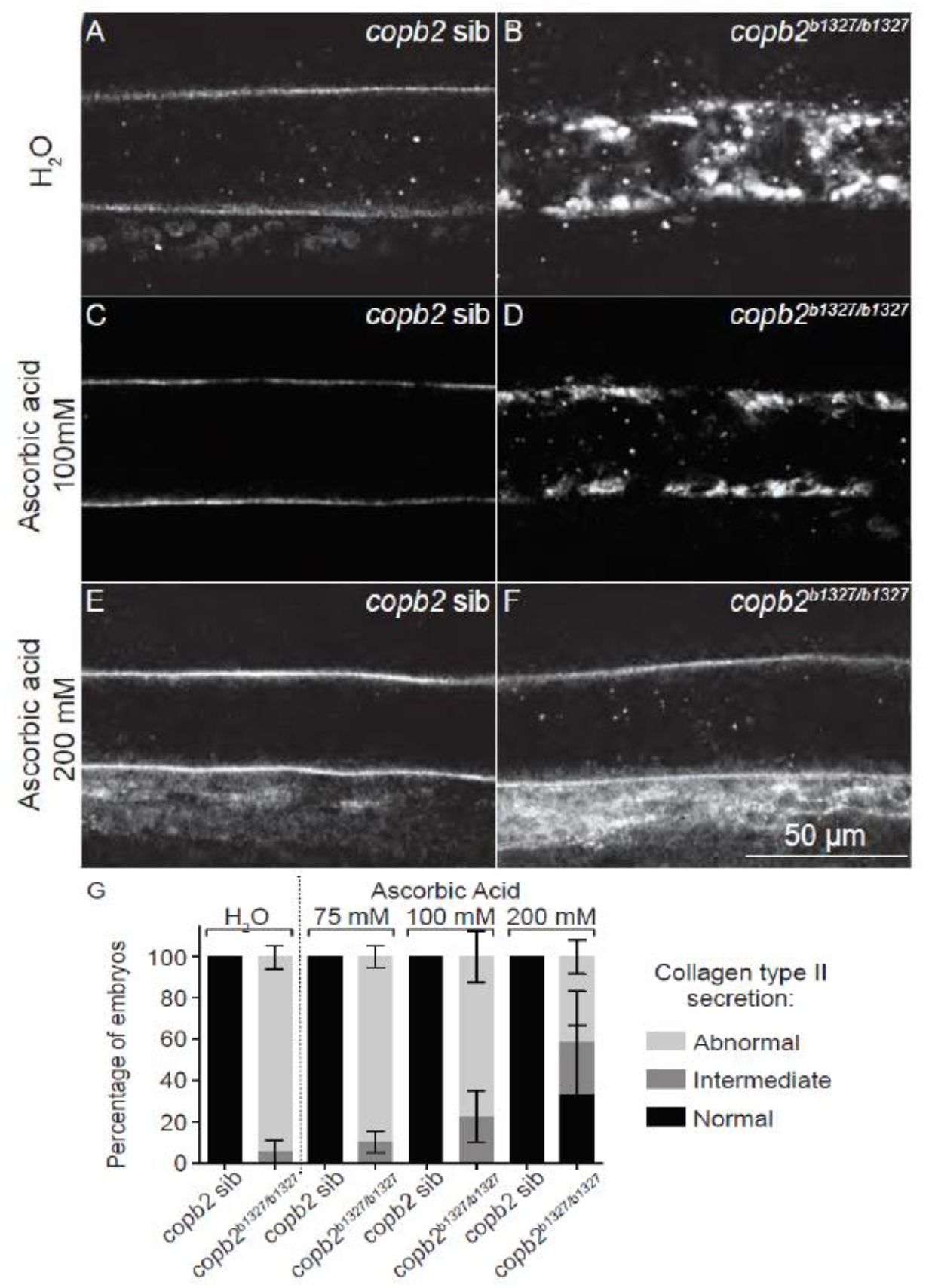
Secretion of collagen type II in *copb2*^*b1327/b1327*^ zebrafish mutants can be rescued by treatment with ascorbic acid. (**A-F)** Confocal sections of zebrafish notochords. Immunolabeling of type II collagen in untreated (A) *copb2* siblings and (B) *copb2*^*b1327/b1327*^ embryos, 100 mM ascorbic acid-treated (C) *copb2* siblings and (D) *copb2*^*b1327/b1327*^ embryos, and 200 mM ascorbic acid-treated (E) *copb2* siblings and (F) *copb2*^*b1327/b1327*^ embryos. **(G)** Distribution of the phenotypes for each condition. Normal indicates wild-type-like phenotype with normal trafficking and secretion of type II collagen, intermediate describes decreased secretion of type II collagen, and abnormal describes mutant-like phenotype with defective trafficking and secretion of type II collagen. Anterior is to the left and dorsal to the top. Images were taken at the level of the yolk extension. Untreated *copb2* siblings, n=10 embryos; untreated *copb2*^*b1327/b1327*^ mutants, n=13 embryos; 75 mM ascorbic acid-treated *copb2* siblings, n=11 embryos; 75 mM ascorbic acid-treated *copb2*^*b1327/b1327*^ mutant, n=22 embryos; 100 mM ascorbic acid-treated *copb2* siblings, n=10 embryos; 100 mM ascorbic acid-treated *copb2*^*b1327/b1327*^ mutant, n=21 embryos; 200 mM ascorbic acid-treated *copb2* siblings, n=5 embryos; 200 mM ascorbic acid-treated *copb2*^*b1327/b1327*^ mutant, n=5 embryos. 30 hpf.

## Discussion

Coatomer complexes and their interacting proteins play an essential role in the anterograde trafficking of cargo lipids and proteins, retrograde transport of proteins from the Golgi complex to ER, maintaining the structure of the Golgi complex and preventing ER and Golgi stress (4, 17, 28, 29). As such, their function is critical in development. Defects in different subunits have been implicated in both animal models and human genetic disorders (2, 4, 5). Here, we report a coatopathy due to pathogenic variants in *COPB2* in five unrelated families. The Subjects in this cohort present with osteopenia and fractures, thus expanding the phenotypic spectrum of previously reported neurodevelopmental features (22). Our data support a role for COPB2 in the intracellular trafficking of collagen as a mechanism for the early-onset osteoporosis.

Type I collagen is the major component of bone extracellular matrix. The biosynthesis of collagen involves multiple steps of post-translational modifications, including hydroxylation of lysine and proline residues in the ER and glycosylation in the Golgi complex (7, 27). Abnormal intracellular trafficking of PC may result in altered post-translational modifications, abnormal folding of the triple helix, and ultimately instability of collagen fibers. Therefore, several defects in vesicular components associated with intracellular collagen trafficking have been reported to cause severe skeletal dysplasias (11–14, 21). The export of fibrillar PC, such as PCI and PCII, is a challenging task for the ER that requires a dedicated set of proteins and, additionally, the retrograde input of membranes from the Golgi/ERGIC (3, 8–10). The coatomer complex COPI is essential for Golgi/ERGIC to ER trafficking. Our results demonstrate that *COPB2* deficiency selectively affects the export of PCI in fibroblasts, providing evidence for the physiological relevance of COPI-mediated retrograde trafficking in the exit of PC from the ER. Curiously, despite the delayed transport causing retention of PC within the ER, mass spectrometry analysis in mouse bones did not detect significant changes in lysine and proline hydroxylation. This suggests that although COPB2 deficiency delays secretion of type I collagen, it does not appear to affect its biosynthesis or post-translational processing in the ER. Dysfunction of the Golgi may result in altered protein glycosylation, as had been reported for defects in other components of the COPI complex (20), and this may be further contributing to the bone fragility.

Previous studies in *Copb2*^*R254C/R254C*^ mice that are homozygous for the p.Arg254Cys variant (Family 5, Subjects 5 and 6) showed normal growth and no obvious brain abnormalities, as opposed to reduced brain size and cortical malformations in the *Copb2*^*R254C/Zfn*^ mouse model (22). Interestingly, neurospheres derived from *Copb2*^*R254C/Zfn*^ and *Copb2*^*R254C/wt*^ mouse models showed significantly reduced cell viability, suggesting that the p.Arg254Cys is a hypomorphic variant (22). Although the molecular mechanism of the neurodevelopmental phenotype in our subjects is not well understood, this study supports a role for *COPB2* in brain development. These findings are in agreement with the existence of other vesicular trafficking defects that are characterized by microcephaly and developmental delay (2, 4).

The anatomical and functional abnormalities seen in both heterozygous *Copb2* mice and homozygous *copb2* zebrafish were at least partially corrected by ascorbic acid supplementation. Ascorbic acid is known to stimulate collagen synthesis and secretion (16, 27), enhance healing of fractured bones in preclinical studies (30), and significantly decrease the risk of osteoporosis and hip fracture in adults (31, 32). Because mice differ from humans and fish by being able to synthesize *de novo* vitamin C, supplementation did increase bone mass but not in a statistically significant fashion. Together, these experiments support the use of ascorbic acid as a therapeutic modality to improve bone fragility in individuals with *COPB2* deficiency and provide a mechanistic rationale for its potential positive effects. Testing the utility of ascorbic acid supplementation in *COPB2*-deficient subjects was beyond the scope and timeframe of this work; however, this remains an important question to address in future studies.

One limitation of our study is the diversity of phenotypes noted within a small cohort of patients. Although our subjects had overlapping skeletal and neurological features, the severity was variable. Moreover, the neurological disability that is more severe in Subjects 2, 5 and 6 likely had contributed to the osteopenia, as would be frequently seen in individuals affected with neuromuscular disorders. Yet, it is important to note that other subjects who present with milder neurological phenotype, such as Subjects 1 and 3 still showed significant osteopenia and fractures (and this skeletal phenotype was recapitulated in the animal models despite not showing any obvious neurological abnormalities). It is also noteworthy that 4/6 subjects had a heterozygous, *de novo* loss-of-function variants while in one family a homozygous missense presumably hypomorphic variant was detected. Given the early embryonic lethality that was observed in 2 independent mouse models in this study, in a previously published mouse model (22), and in zebrafish, complete loss of function of *COPB2* is likely not tolerated in humans. The probability of loss of function intolerance (pLI) score reported in gnomAD (23) is 1 and supports this prediction. It is difficult to discuss genotype-phenotype correlation in such a small cohort, and additional individuals will need to be identified before the spectrum of *COPB2*-related disorder could be fully characterized.

In summary, we have identified *COPB2* loss-of-function variants as a cause of coatopathy with developmental delay and juvenile osteoporosis. Early-onset osteoporosis increases the risk for osteopenia and fractures in adulthood. Identification of individuals with low BMD early in life may enable preventive intervention to reduce the risk of fractures and adult osteoporosis. Our findings highlight the important role of the COPI complex and *COPB2* in bone development and demonstrate that collagen trafficking is a biological determinant of bone mass, independent of bone quality that is affected in OI. Our work on this rare disorder suggests that decreased collagen flux may contribute to more common forms of osteoporosis.

## Materials and Methods

### Patients

Prior to research studies, informed consent was obtained as part of the Undiagnosed Diseases Network for Subjects 1 and 2, according to the institutional review board of the National Human Genome Research Institute. The research sequencing study (exome and genome sequencing) and transfer of clinical and sequencing data were approved by the human subject ethics committees as part of the 100,000 Genomes Project for Subject 3, and at Baylor College of Medicine for Subjects 4, 5 and 6. All families provided informed consent prior to participation.

### Exome or genome sequencing and identification of C*OPB2* variants

**Subject 1** had research trio exome sequencing performed at the Baylor College of Medicine Human Genome Sequencing Center (BCM-HGSC). Briefly, library was constructed as described in the BCM-HGSC protocol (https://www.hgsc.bcm.edu/content/protocols-sequencing-library-construction) and hybridized to the HGSC VCRome 2.1 design (42Mb NimbleGen, Cat. No. 06266380001) (33). Sequencing was performed in paired-end mode using the Illumina HiSeq 2000 platform, with sequencing-by-synthesis reactions extended for 101 cycles from each end and an additional 7 cycles for the index read. With a sequencing yield of 6.6 Gb, the samples achieved 92% of the targeted exome bases covered to a depth of 20X or greater. Illumina sequence analysis was performed using the HGSC Mercury analysis pipeline (34, 35) (https://www.hgsc.bcm.edu/software/mercury) which moves data through various analysis tools from the initial sequence generation on the instrument to annotated variant calls. In parallel to the exome workflow an Illumina Infinium Exome-24 v1.1 array was generated for a final quality assessment. This included orthogonal confirmation of sample identity and purity using the Error Rate In Sequencing (ERIS) pipeline developed at the HGSC. A successfully sequenced samples met quality control metrics of ERIS SNP array concordance (>90%) and ERIS average contamination rate (<5%).

GeneMatcher (https://genematcher.org/)(36) assisted in the recruitment of **Subject 2**, who had clinical exome sequencing at GeneDx. **Subject 3** had genome sequencing through the UK 100,000 Genomes Project. In brief, DNA was extracted from lymphocytes at the North Thames Genomic Laboratory Hub, Great Ormond Street Hospital for Children NHS Foundation Trust, and sent to Illumina for genome sequencing. Sequencing data was passed through Genomics England's bioinformatics pipeline for alignment, annotation and variant calling. Based on the Human Phenotype Ontology terms entered for this patient, the following gene panel was applied: Osteogenesis Imperfecta 1.13 (panelapp.genomicsengland.co.uk). The *COPB2* c.1206-2A>G variant was not called by the PanelApp but was identified subsequently using Exomiser tool (37). The variant was confirmed via bidirectional sanger sequencing at the North Thames Genomic Laboratory Hub, Great Ormond Street Hospital for Children, NHS Foundation Trust. **Subject 4** had clinical sequencing at GeneDx. The sequencing analysis for **Subjects 5 and 6** was reported in a prior publication (22).

### RNA sequencing

Messenger RNA from whole blood was extracted, quantified, and processed, prior to library preparation using the Illumina TruSeq kit (Illumina, SD) according to manufacturer’s instructions. Samples were subsequently multiplexed and subject to 150bp paired-end sequencing on the Illumina HiSeq2000 platform. Resulting raw bcl reads were converted to FASTQ prior to being processed using a pipeline adapted from the GTEX Consortium (38). Briefly, reads were aligned to the GRCh37 human reference using STAR (39), followed by gene-based quantification with RSEM (40) and alignment visualization in IGV (41).

### RNA extraction and real-time qPCR

Total RNA was extracted from lymphoblastoid cells derived from Subject 1, her parents and sibling using Trizol reagent. Similarly, total RNA was extracted from fibroblasts derived from Subject 2 and her mother. The Superscript III First Strand RT‐PCR kit was used to synthesize cDNA according to the manufacturer's protocol (Invitrogen). Real‐time qPCR was performed on LightCycler instrument (Roche) with *GAPDH* as internal control.

### Cell culture and PC transport assay

All cells were cultured at 37°C and 5% CO2 in a humid environment. HeLa (ATCC) and BJ-5ta hTERT-immortalized fibroblasts (ATCC), referred to as human fibroblasts (HFs) were cultured according to ATCC guidelines. The following *COPB2* siRNA sequences were used in this study: 1) CGAUGUAUCUCCUAGGCUA; 2) GGUCAAACAAUGUCGCUUU and 3) CAGUAUCCACAGAUCCUGA. HeLa and HFs cells were treated for 36 hours with Oligofectamine (Life Technologies) or RNAiMax (Life Technologies), respectively, using different final concentrations of siRNAs (20 nM and 100 nM) to obtain different levels of silencing. Total cell lysates were analyzed by Western blot to quantify the COPB2 levels as described previously (12). Primary antibodies used in this study were: mouse monoclonal anti-coatomer (CM1A10 (42)), anti-GM130 (BD Transduction), anti-Giantin (Abcam), anti-GBF1 (BD Transduction), anti-KDEL receptor (Enzo LifeSciences), sheep polyclonal anti-TGN46 (SeroTech), rabbit polyclonal anti-PC-I (Rockland), anti-VSV-G (Bethyl), anti-COPB2 (Thermo Scientific), and anti-ß-actin (Sigma). Immunofluorescence experiments were performed as previously described (12). All the images were acquired using a Zeiss confocal microscope LSM800. All the transport assays were performed as previously described (12). Briefly, to follow endogenous PC-I in fibroblasts, cells were incubated for 3 hours at 40°C in DMEM supplemented with 1% serum and 20 mM HEPES pH 7.2, then shifted to 32°C in the presence of cycloheximide (100 μg/ml) and ascorbate (50 μg/ml) for the indicated times. Similarly, for VSV-G assays, cells were infected with the VSV-ts045 variant for 1 hour at 32°C in serum-free medium. After infection, the cells were washed several times and shifted for 3 hours at 40°C in complete medium. Cycloheximide was added to a final concentration of 100 μg/ml and the cells were shifted to 32°C for the indicated times.

Quantification of GBF1-positive and COPI-positive structures was performed using Fiji software with the same threshold for all the analyzed conditions. 50 cells/condition from three independent experiments were counted. Quantitative evaluation of PC-I and VSV-G transport was performed by analyzing the immunofluorescence staining patterns of at least 100 cells in three independent experiments.

### Zebrafish CRISPR/Cas9 model

#### Generation of mutant lines using CRISPR/Cas9

One single guide RNA (sgRNA) was designed that contained targeting sequences in exon 2 (5’-GGACATCAAGCGCAAACTCA-3’) of *copb2*. sgRNA and cas9 RNAs were co-injected at the 1-cell stage. The following primers: forward 5’-AGAGTACTGATGTATAATTTCTGC-3’ and reverse: 5’-TCACTTGTGTTTCATGGTTC-3’ were used to identify introduced mutations. Wild-type ABCxTu and heterozygous mutant copb2^b1327/+^ adult zebrafish were maintained as previously described (43). Embryos and larvae were staged according to the staging series (44) or by hours postfertilization (hpf) or days postfertilization (dpf). Siblings (sibs) are defined as a mix of homozygous wild-type (WT) and heterozygous mutants from intercrosses of heterozygous mutant adults. All experimental procedures were approved by the University of Oregon IACUC.

#### Immunolabeling, Alcian blue/Alizarin red staining

Labeling of whole-mount larvae was performed as previously described (45) with minor modifications. Thirty hpf embryos were fixed in BT fix (43) overnight and permeabilized with proteinase K (10 μg/ml) for 10 minutes. The following antibodies were used: mouse anti-KDEL (Calbiochem, 1:500), mouse anti-GM-130 (BD Transduction Laboratories, 1:500), mouse anti-collagen type II (Developmental Studies Hybridoma Bank, II-II6B3; 1:200), and biotinylated horse anti-mouse (Vector Laboratories, 1:500). Alcian blue/Alizarin red staining was performed on 7 dpf larvae as previously described (46). Images of larvae stained with Alcian blue/alizarin red were acquired using a Zeiss Axioplan2 compound microscope. Images of alizarin red stained opercula and immunolabeled embryos were acquired using a Zeiss LSM 5 confocal microscope and analyzed using ImageJ.

#### Ascorbic acid treatment

Homozygous *copb2*^*b1327/b1327*^ embryos were treated from 5 hpf, when the notochord has not yet formed, to 30 hpf with 75 mM, 100 mM, and 200 mM ascorbic acid (Sigma). Ascorbic acid was diluted in embryo medium, and the pH of the solution was adjusted to 6.5. Control larvae were treated with an equal volume of water also diluted in embryo medium. At 30 hpf each embryo was scored for its overall morphology and bisected at the end of the yolk extension. The posterior part was used for genotyping and the anterior part was fixed for immunolabeling. After genotyping and immunolabeling, the notochord was imaged.

#### *In situ* hybridization and histology

Assay was performed as previously described (47). Larvae were hybridized with a digoxigenin labeled RNA probe spanning an 812 bp coding sequence between exons 8 and 14 of *copb2*. Stained larvae were embedded in 1 % agarose, 0.5 % agar, 5 % sucrose medium and 16 μm cryosections were cut. Images were acquired using a Zeiss Axioplan2 compound microscope.

### Mouse CRISPR/Cas9 model

We generated a knockout mouse model using CRISPR/Cas9 technology, by introducing a deletion within the coding region of mouse *Copb2*. Guide RNAs (sgRNA) were designed that contained targeting sequences to allow for exon 6 and 7 deletion [5’-CGGGTTTACCTGCCCAGCGG(TGG)-3’ and 5’-ACAGACCTCGGTTCAAAATC(AGG)-3’] of *Copb2*. sgRNA and cas9 mRNA were co-injected into the cytoplasm of C57BL/6J mouse embryos at the 1-cell stage. PCR cloning followed by Sanger sequencing, using primers: 5’-TCCAAGCATTATCCAAGGAAGT-3’, 5’-AACACCAGAGCCAAGAAGTG-3’ and 5’-GCCTTTTTCATGTCCTTCCA-3’, confirmed deletion of exon 6 through exon 7 in founder animals (**Supplemental Figure 2**). Founders were backcrossed to generate N1 animals, inheritance of the deletion in N1 animals was confirmed by Sanger sequencing, and the N1 generation was backcrossed to generate N2 mice. N2 animals were intercrossed to generate the mouse line eventually used for phenotyping. Upon further breeding, no homozygous mice were detected, likely due to early embryonic lethality of the *Copb2*^−/−^ pups. Phenotypic analysis was therefore performed using mice that are heterozygous for the *Copb2* deletion or their wild-type littermates.

In parallel, the BCM component of KOMP2 generated a separate knockout mouse model of *Copb2* (*Copb2*^*em1(IMPC)Bay*^) on the C57BL6/NJ background using the same sgRNAs targeting exon 6. Adult and embryo phenotyping data for this line is available on the IMPC website (https://www.mousephenotype.org).

Mice were housed in the Baylor College of Medicine Animal Vivarium, and all studies were approved by the Institutional Animal Care and Use Committee (IACUC) at the Baylor College of Medicine. *Copb2*^*+/−*^ mice and wild-type littermates were euthanized at 8 weeks of age. For the micro-CT analysis, both males and females were studied. For other studies, only females were studied since bone mineral density was appreciated to be more strongly affected in females. The investigators were blinded to genotype in the micro‐CT, biomechanical, and collagen mass-spectrometry analyses.

#### Micro-CT analysis

Spines and left femurs were scanned in 70% ethanol using a Scanco μCT-40 micro-CT system (55kVp and 145μA X-ray source), and scans were reconstructed at a 16μm isotropic voxel size. Trabecular bone of L4 vertebrae and the distal metaphyses of left femurs were analyzed using Scanco software by manually contouring trabecular bone. For vertebrae, the region of interest (ROI) was defined as the trabecular volume between the L4 vertebral endplates. At femoral trabecular ROI 75 slides (=1.2 mm) were analyzed proximal to the distal femoral growth plate. Quantification of trabecular parameters was performed using the Scanco software with a threshold value of 210. These parameters include bone volume/total volume (BV/TV), trabecular number (Tb.N), trabecular thickness (Tb.Th), connectivity density (Conn.D) and tissue mineral density (TMD) (48). Femur length was measured from the top of the femoral head to the bottom of the medial condoyle. Cortical bone parameters of the femoral midshaft were measured at the exact center and at the distal 75% of femur length using the automated thresholding algorithm included in the Scanco software. Trabeculae in contact with cortical bone were manually removed from the ROI (11 slides analyzed per location, threshold 210). The cortical parameters include total cross-sectional area (Tt.Ar), cortical bone area (Ct.Ar), marrow area (Ma.Ar), cortical thickness (Ct.Th), cross-sectional moments of inertia (CSMI), anterior-posterior diameter, and tissue mineral density (TMD) (48). Ten spines and femurs were scanned per group.

#### Biomechanical measurement

At 8 weeks of age, femurs were collected from euthanized mice, stripped of soft tissues, wrapped in saline‐soaked gauze, and frozen at −20°C until analyses were performed. The femurs were tested to failure in three-point bending at a rate of 0.05mm/sec and were oriented in the test fixture such that the anterior surface was in compression and the posterior surface in tension. Femurs were tested wet at room temperature using displacement mode with an Instron 5848 microtester (Instron Inc., Norwood MA). The test fixture span was adjusted according to the femur length and varied between 6.72 and 7.32mm. A 100N load cell was used to collect data, and load and displacement data were captured at rate of 40Hz by using BLUEHILL Software (Instron 5848). The maximum load was determined by finding the highest load value recorded before the specimen fractured. The region of the load-displacement curve between 1N and the maximum load was separated into 5 segments and the fitted line of the segment with greatest slope was defined as the stiffness. A line representing 10% degradation of this stiffness was used to define the yield point. The elastic region was identified as the region from the completion of the preload to the yield point. The post-yield region was identified as the region from the yield point until point of specimen fracture. Using a trapezoidal numerical integration method, the work to fracture was calculated as the area under the load-displacement curve. Since different span lengths were used according to the length of each tested femur, the mechanical properties were adjusted to account for the test-fixture geometry (49). The adjusted parameters are maximum bending moment, adjusted post-yield displacement, rigidity, and adjusted work-to-failure. The cross-sectional geometry of each bone as determined by micro-CT image analysis was used to convert the maximum load and stiffness data into ultimate stress and elastic modulus values using beam theory.

#### Bone histomorphometry

Calcein (250 μg i.p.) was injected 7 days prior to euthanasia, and alizarin red (800 μg i.p.) was injected 2 days before euthanasia. Spines were collected and fixed in 10% formalin for 48 hours. Mouse spine samples were then embedded in plastic and sectioned with tungsten carbide blades. Trichrome staining and tartrate-resistant acid phosphatase staining were performed for visualizing osteoblasts and osteoclasts, respectively. Calcein and alizarin red double labeling was used to study dynamic histomorphometry. Results were quantified with the Bioquant Osteo 2014 Image Analysis System for evaluation of bone formation and resorption parameters.

#### Mass spectrometry of type I collagen

Collagen was prepared from minced femurs, collected from euthanized 8-week-old mice. Type I α-chains were extracted by heat denaturation (90°C) in SDS-PAGE sample buffer, resolved on 6% SDS-PAGE gels (50), cut from gels and digested with trypsin in-gel (51). Bone tissue was also digested with bacterial collagenase as described (52). Collagenase-generated peptides were separated by reverse phase HPLC (C8, Brownlee Aquapore RP-300, 4.6 mm × 25 cm) with a linear gradient of acetonitrile:n-propanol (3:1 v/v) in aqueous 0.1% (v/v) trifluoroacetic acid (53). Individual fractions were analyzed by Liquid Chromatography/Mass spectrometry (LC/MS). Peptides were analyzed by electrospray LC/MS using an LTQ XL ion-trap mass spectrometer (Thermo Scientific) equipped with in-line liquid chromatography on a C4 5μm capillary column (300 μm × 150 mm; Higgins Analytical RS-15M3-W045) and eluted at 4.5 μl/min. The LC mobile phase consisted of buffer A (0.1% formic acid in MilliQ water) and buffer B (0.1% formic acid in 3:1 acetonitrile:n-propanol v/v). An electrospray ionization source (ESI) introduced the LC sample stream into the mass spectrometer with a spray voltage of 3kV. Proteome Discoverer search software (Thermo Scientific) was used for peptide identification using the NCBI protein database.

#### Ascorbic acid treatment

*Copb2*^*+/−*^ female mice were provided with ascorbic acid-enriched chow or the control chow for a period of 5 weeks. The ascorbic acid-enriched chow (ENVIGO Teklad laboratory animal diet) contains 18% protein, 49.1% carbohydrate and 6.3% fat, in addition to 1% ascorbic acid.

The control chow (ENVIGO Teklad laboratory animal diet) contains 18.2% protein, 49.6% carbohydrate and 6.4% fat, without ascorbic acid. Experimental diet was initiated at weaning age (P21) and mice were collected for analysis by μCT of spines at 8 weeks of age.

### Statistical analysis

Nonparametric two-tailed paired t test or one-way ANOVA were used as indicated to compare between samples. Statistical significance was determined as p value that is equal to, or less than 0.05.

## Author Contributions

BL, MADM, MW, RM, LCB, AC, BBS, and RV designed the study, performed the experiments and wrote the draft of this manuscript. MJ, DAS, JAR, ZJ, SC, VRS, MS, GM, CDV, RR, ADC, AS, TTM, JA, RWS, LTE, DRM, RAG, NAH, GEZ, NRM, DMM and SNJ contributed to the recruitment of the subjects and their families, their clinical phenotyping, genomic sequencing and RNA studies. IG, DGL, XL, KSJ, YCL, IWS, JMS, DB, BCD, YC, MMJ, EM, CK, MAW, MED, JRS, LDH, DRE and CGA participated in the generation and phenotyping of the mouse model. BAT, JBP and JW contributed to the generation and phenotyping of the zebrafish model. All authors reviewed and edited the manuscript. This study was conducted as part of the Undiagnosed Diseases Network (UDN), and a full list of members of the UDN is available under Supplementary Materials. A full list of members of the Genomics England Research Consortium is available under Supplementary Materials.

## Acknowledgments

The authors would like to thank the patients and their families for participation in the study. We thank Paula Patricia Hernandez, Judy Peirce, and Poh Kheng Loi for their technical assistance. RM was supported by the Michael Geisman-Osteogenesis Imperfecta Foundation (OIF) Fellowship Award, the T32GM007526-41 and the Lawrence Bone Disease Program of Texas Research Award. This work was also supported by NIH U01HG007709 (BL), NIH P01 HD070394 (BL), NIH U54 AR068069 (BL, VRS), NIH UM1HG006348 (AB, MD, JDH), NIH U54NS093793 (MW), NIH K08 DK106453 (LCB), NIH/NIGMS T32GM007526 (MJ), NIH/NIDCR R03DE026233 (IG), NIH 5UM1HG006542 (VRS), NIH/NIAMS R37 AR037318 (DRE) and NIH/NICHD U54HD083092 for the Baylor College of Medicine Intellectual and Developmental Disabilities Research Center (IDDRC). LCB was also supported by a Career Award for Medical Scientists from the Burroughs Wellcome Fund. MADM acknowledges the support of Telethon grant TGM11CB1, Associazione Italiana per la Ricerca sul Cancro grant IG2013_14761, and European Research Council Advanced Investigator grant 670881 (SYS MET). GMM was supported by the National Institute of Neurological Disorders and Stroke (NINDS) under award number K08NS092898, Jordan’s Guardian Angels and the Brotman Baty Institute. This work was also supported by grant #3092-51000-057-04S from the US Department of Agriculture to the LTG core at Children’s Nutrition Research Center at Baylor College of Medicine. The content is solely the responsibility of the authors and does not necessarily represent the official views of the National Institutes of Health. See supplemental acknowledgments for consortium details (Undiagnosed Diseases Network and the Genomics England Research Consortium).

