## Supplemental data for "*COPB2* haploinsufficiency causes a coatopathy with osteoporosis and developmental delay"

### Supplementary Appendix

**Supplemental Figure 1.** *COPB2* variant c.1237_1238delAA (p.Lys413Aspfs*3) leads to nonsense mediated decay (NMD) in Subject 1.

**Supplemental Figure 2.** Generation of *Copb2* null allele by CRISPR/Cas9 in mice.

**Supplemental Figure 3.** *Copb2*$^{+/-}$ mice do not show significant alterations in bone formation rate and mineralization.

**Supplemental Figure 4.** Collagen expression and post-translational modification is not altered in *Copb2*$^{+/-}$ mice.

**Supplemental Figure 5.** Generation of *copb2* mutant allele by CRISPR/Cas9 in zebrafish.

**Supplemental Figure 6.** No obvious brain abnormalities were observed in *copb2*$^{b1327/b1327}$ mutants.

**Supplemental Figure 7.** Partial depletion of COPB2 induces disorganization of the Golgi complex but does not impair ER exit of VSVG.

**Supplemental Figure 8.** Notochord cell organization in *copb2*$^{b1327/b1327}$ mutants can be rescued by addition of ascorbic acid.

**Supplemental Figure 9.** Ascorbic acid-enriched diet is associated with trend of improvement in *Copb2*$^{+/-}$ mouse.

**Supplemental Table 1.** Summary of biomechanical testing in Copb2$^{+/-}$ mouse femurs.

**Members of the Undiagnosed Diseases Network**

**Members of the Genomics England Research Consortium**

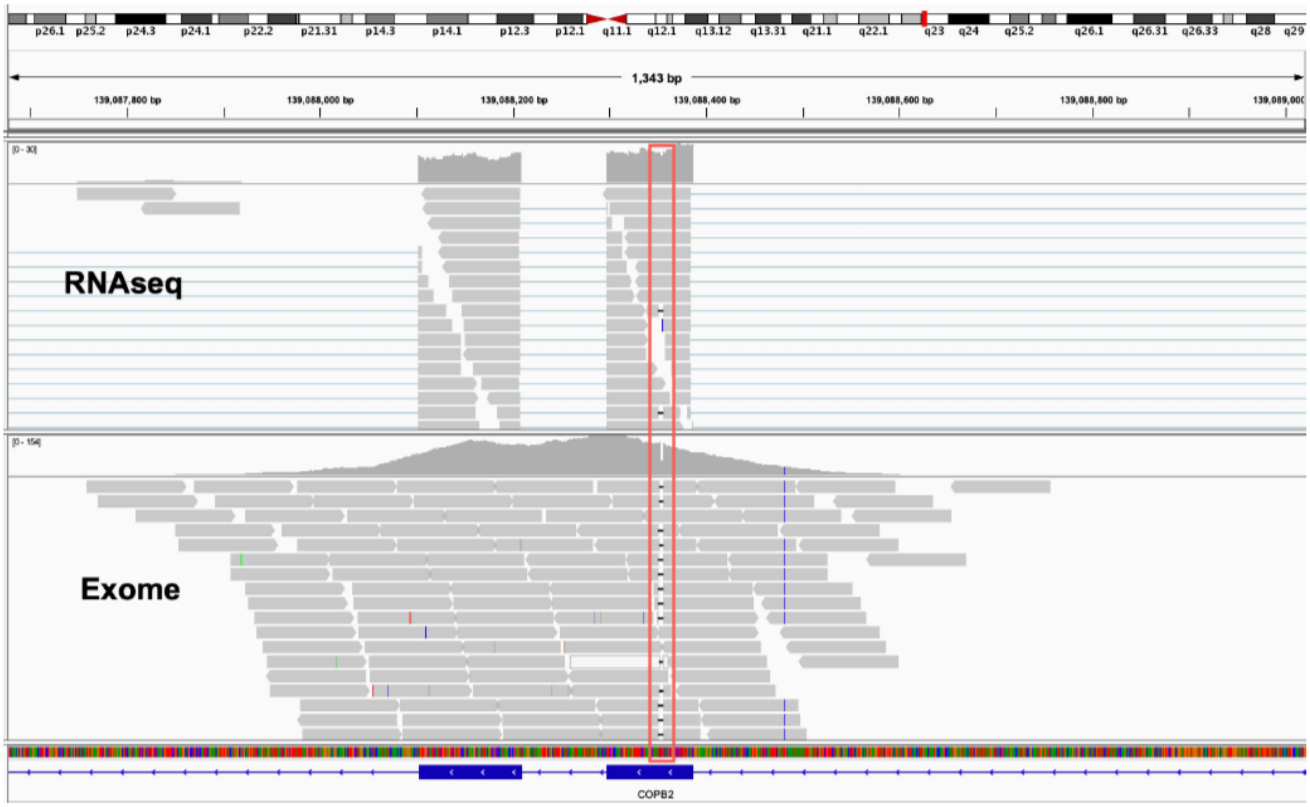


**Supplemental Figure 1. *COPB2* variant c.1237_1238delAA (p.Lys413Aspfs*3) leads to nonsense mediated decay (NMD) in Subject 1.** Integrated Genomic Viewer (IGV) illustration of RNA sequencing (top) and exome sequencing (bottom) data for Subject 1, carrying a c.1237_1238delAA (p.Lys413Aspfs*3) variant in *COPB2*. Exome sequencing detected the variant in 66/122 (54%) reads, consistent with heterozygosity. RNA sequencing detected only 2/20 (10%) reads with a deletion, supporting nonsense-mediated decay.

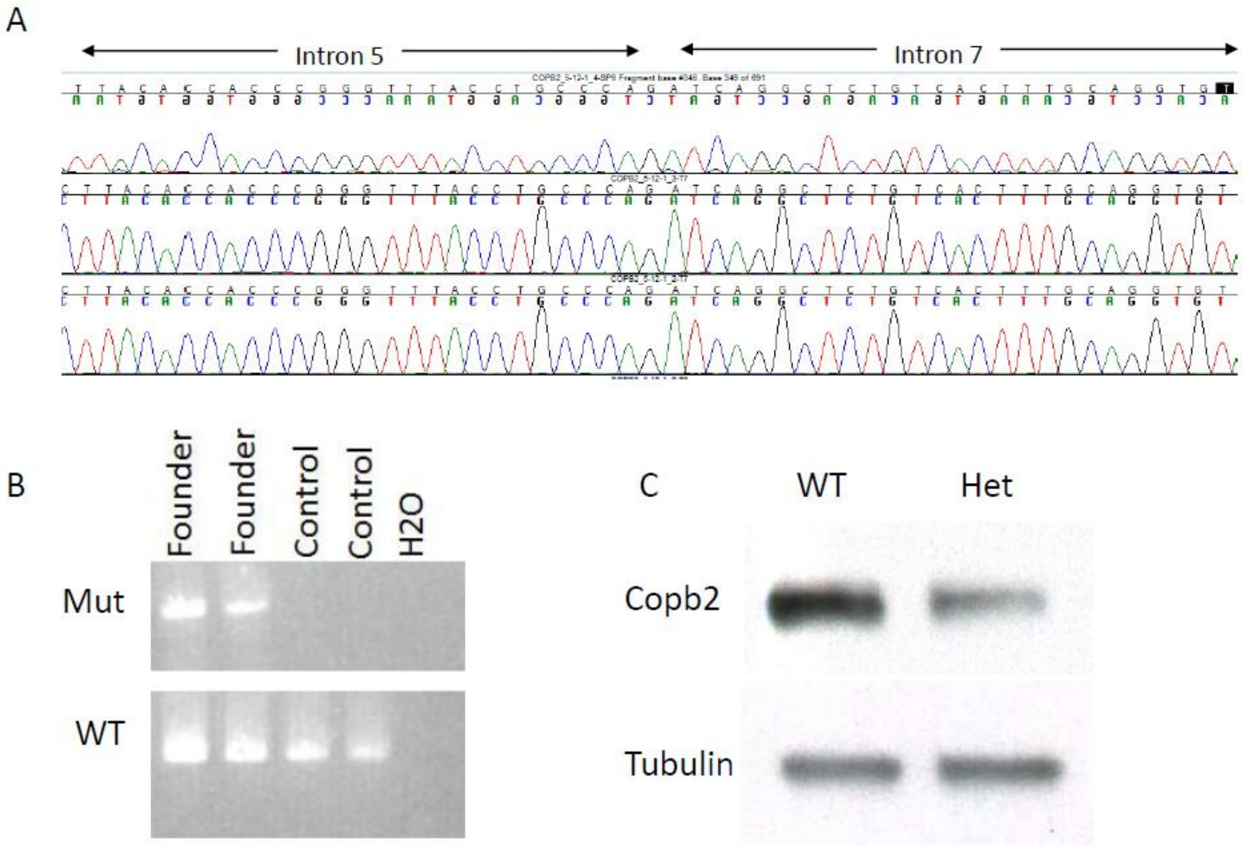


**Supplemental Figure 2. Generation of *Copb2* null allele by CRISPR/Cas9 in mice. (A-B)** Guide RNAs (sgRNA) were designed to allow for deletion of exon 6 through 7 of *Copb2*. Correct targeting with loss of exons 6 and 7 was verified by PCR of genomic DNA and Sanger sequencing. **(C)** Heterozygosity for *Copb2* deletion was associated with Copb2 protein reduction, as shown by western blot analysis in protein extracted from mouse fibroblasts (Het - *Copb2*$^{+/-}$ and WT – wild type littermate).

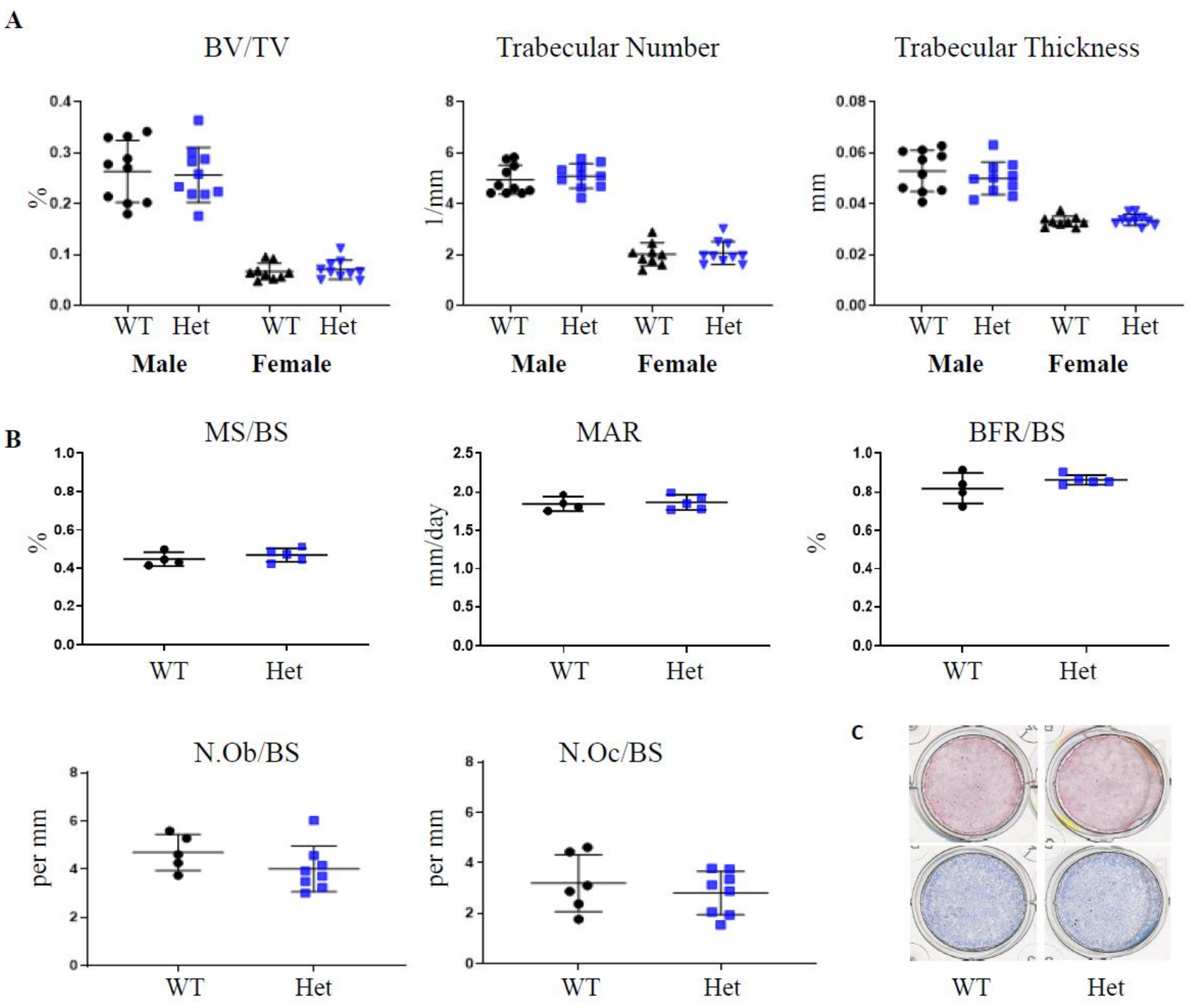


**Supplemental Figure 3.** ***Copb2***$^{+/-}$ **mice do not show significant alterations in bone formation rate and mineralization. (A)** Micro CT analysis of femurs in *Copb2*$^{+/-}$ mice (Het) did not show significant difference in bone architectural parameters compared to wild type littermates (WT). (B) Results of bone histomorphometry study of spine (L4 vertebrae) in *Copb2*$^{+/-}$ female (Het) and wild type littermates (WT) are shown for analyses of mineralized surface/bone surface (MS/BS), mineralized apposition rate (MAR), bone formation rate/bone surface (BFR/BS), osteoblast number/bone surface (N.Ob/BS) and osteoclast number/bone surface (N.Oc/BS). (C) *In vitro* calvaria cultures did not show significant difference in mineralization (alizarin red staining, upper panel) or in alkaline phosphatase activity (lower panel).

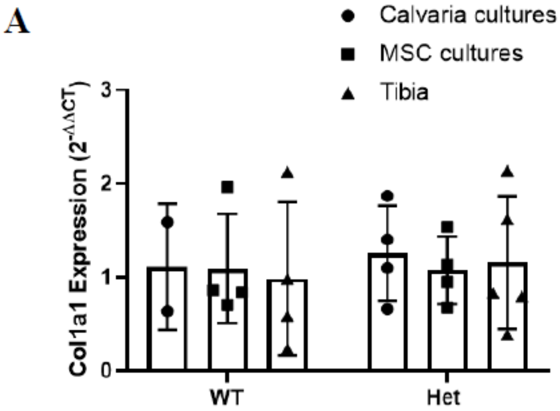


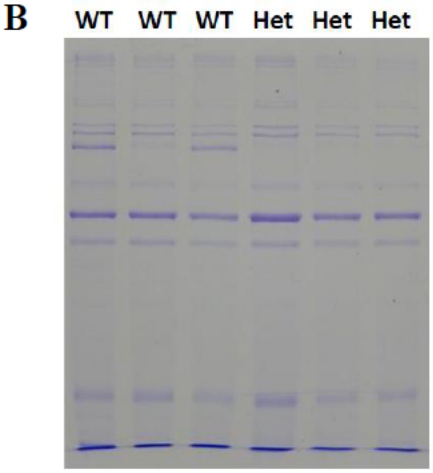


C

| Col1a1(I) | | | | | | | |
|---|---|---|---|---|---|---|---|
| sample | P771 | P876 | P948 | P1011 | K174 | K252 | K756 |
| #81 Het | 50 | 62 | 75 | 34 | 75 | 25 | 40 |
| #71 Het | 49 | 54 | 74 | 42 | 76 | 26 | 42 |
| #72 Het | 42 | 59 | 66 | 41 | 76 | 25 | 52 |
| #84 WT | 59 | 63 | 79 | 59 | 71 | 25 | 39 |
| #70 WT | 46 | 58 | 81 | 53 | 71 | 27 | 39 |
| #83 WT | 56 | 59 | 69 | 52 | 68 | 26 | 32 |

| Col1a2(I) | | | | | | |
|---|---|---|---|---|---|---|
| sample | P81 | P486 | P504 | P804 | P707 3Hyp | P891 |
| #81 Het | 77 | 90 | 90 | 86 | 12 | 84 |
| #71 Het | 79 | 89 | 83 | 88 | 20 | 81 |
| #72 Het | 74 | 90 | 89 | 91 | 14 | 85 |
| #84 WT | 83 | 92 | 88 | 86 | 17 | 83 |
| #70 WT | 81 | 92 | 86 | 83 | 19 | 84 |
| #83 WT | 79 | 86 | 78 | 88 | 17 | 81 |

**Supplemental Figure 4. Collagen expression and post-translational modification is not altered in *Copb2*$^{+/-}$ mice. (A)** Quantitative PCR analysis of *Col1a1* expression in cDNA derived from calvaria cultures, marrow stromal cells culture (MSC) and tibia, results normalized to *B2m* (independent values represent biological repeats, n=2-5 per sample). **(B)** Type I collagen isolated from femurs of 8 week old mice showed no difference in electrophoretic migration pattern between *Copb2*$^{+/-}$ (Het) and wild type littermates (WT). **(C)** There are no changes in the hydroxylation pattern of proline and lysine residues. Results indicate measured % hydroxylation at the indicated residue sites.

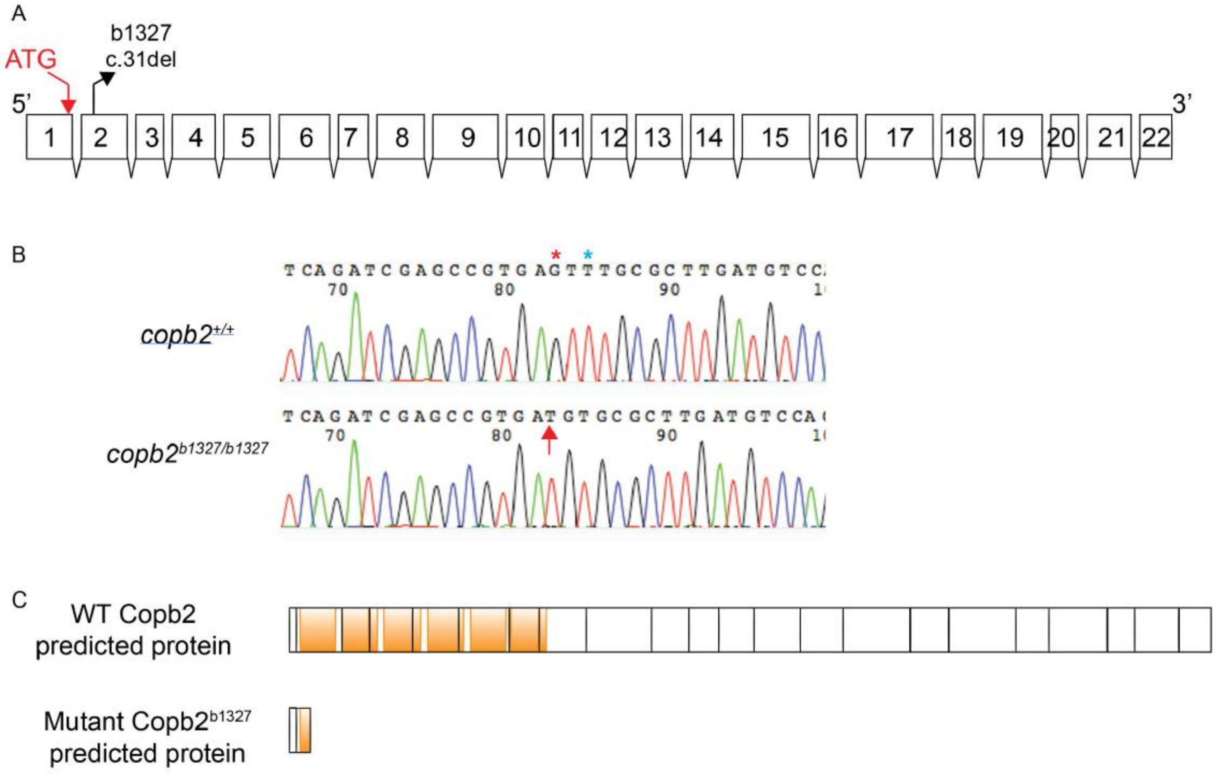


**Supplemental Figure 5. Generation of *copb2* mutant allele by CRISPR/Cas9 in zebrafish.**

**(A)** Nature and localization of *copb2* mutations. Mutation in *copb2*$^{b1327}$ is a 1 bp substitution at position 29 from the canonical ATG in the reference sequence (NM_001001940) and a 1 bp deletion at position 31. **(B)** Sequence chromatograms of *copb2*$^{+/+}$ and *copb2*$^{b1327/b1327}$. Red star indicates the nucleotide present in the WT allele and deleted in the *b1327* allele. Blue star indicates the nucleotide present in the WT allele and substituted in the *b1327* allele. Arrow indicates the position of the deletion in the mutant sequence. The reverse-complement sequence is presented. **(C)** Predicted proteins of WT copb2 and mutant copb2$^{b1327}$. Orange boxes represent WD40 domains.

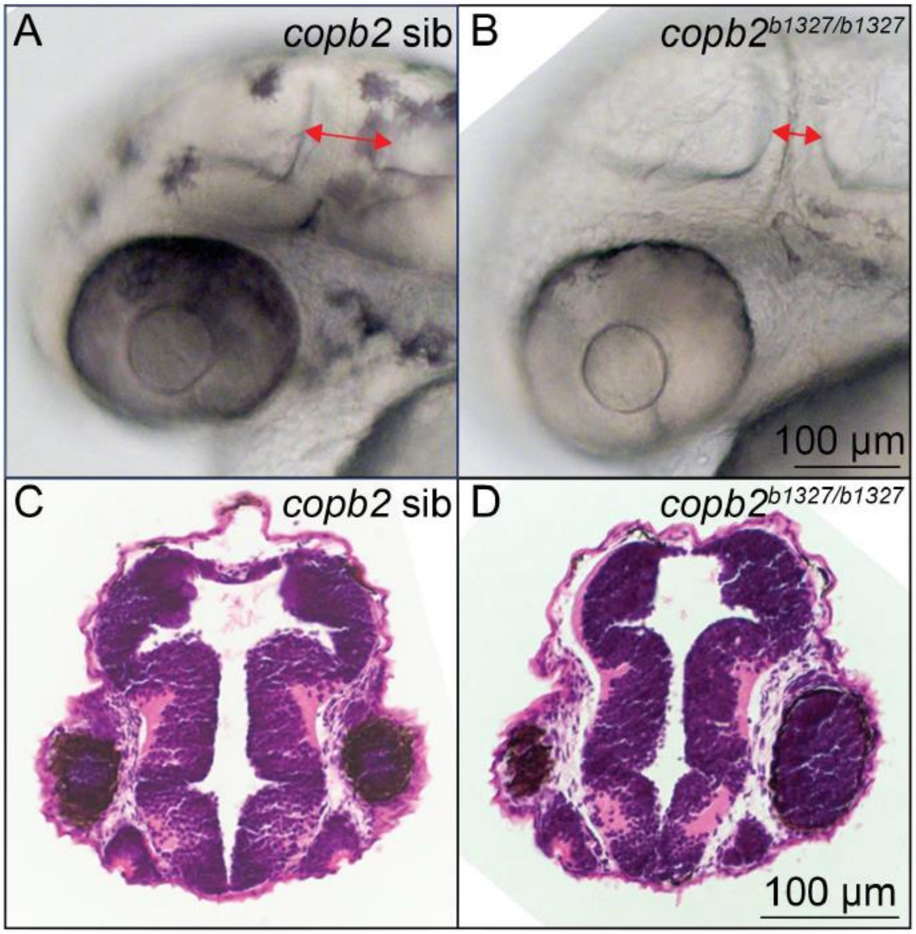


**Supplemental Figure 6. No obvious brain abnormalities were observed in *copb2*$^{b1327/b1327}$ mutants.** Differential interference contrast images of the midbrain-hindbrain boundary in **(A)** *copb2* siblings and **(B)** *copb2*$^{b1327/b1327}$ mutant embryos. Red double arrow indicates the width of the midbrain-hindbrain boundary. Hematoxylin and eosin stain of brain sections from **(C)** *copb2* siblings and **(D)** *copb2*$^{b1327/b1327}$ mutant embryos at 30 hpf.

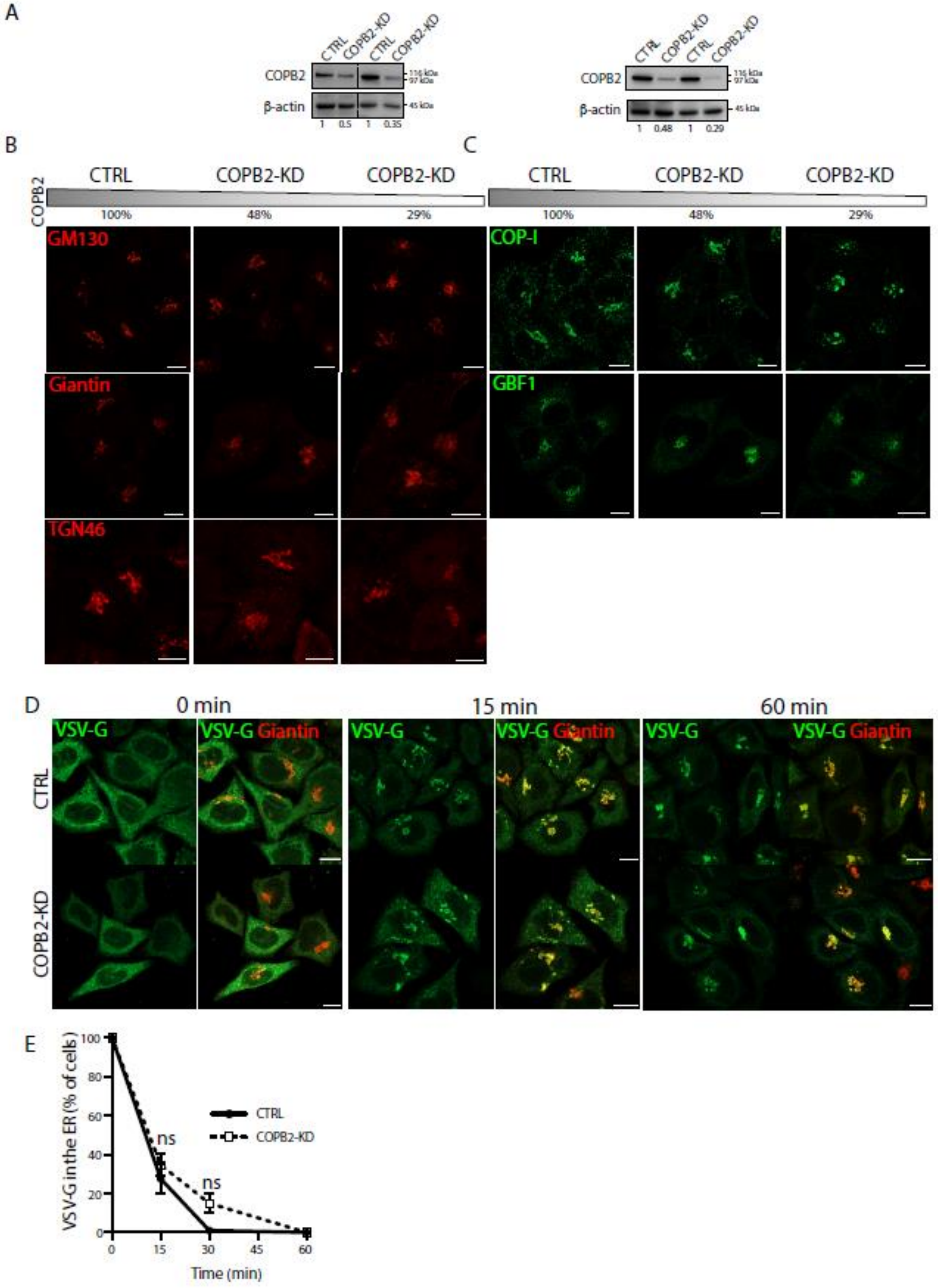
A
CTRL
COPB2-KD
COPB2
β-actin
116 kDa
97 kDa
45 kDa
1 0.5 1 0.35
1 0.48 1 0.29
B
C
CTRL
COPB2-KD
COPB2
100%
48%
29%
GM130
Giantin
TGN46
COP-I
GBF1
D
0 min
15 min
60 min
VSV-G
VSV-G Giantin
CTRL
COPB2-KD
E
VSV-G in the ER (% of cells)
Time (min)
ns
CTRL
COPB2-KD

**Supplemental Figure 7. Partial depletion of COPB2 induces disorganization of the Golgi complex but does not impair ER exit of VSVG. (A)** Western blot analyses of COPB2 in human fibroblats (HF, left) and HeLa (right) cells, mock (CTRL) or COPB2-siRNA treated for 36 hours to obtain different levels of COPB2 reduction (50% and 35% using 20 nM and 100 nM siRNAs, respectively in human fibroblasts, and 48% and 29% using 20 nM and 100 nM siRNAs, respectively in HeLa cells). Numbers represent COPB2 protein levels normalized for beta-actin. **(B)** HeLa cells were mock (CTRL) or COPB2-siRNA treated as in A. The gray bar indicates the residual COPB2 protein level after siRNA treatment. Cells were subjected to indirect immunofluorescence using an antibody against the cis-Golgi marker GM130 (upper panel), medial Golgi marker Giantin (middle panel) and trans-Golgi protein TGN46 (lower panel). Scale bars=10µm. **(C)** Immunofluorescence analyses of CTRL and COPB2-KD HeLa cells for the ER-Golgi intermediate compartment (ERGIC) proteins COPI and GBF1. Scale bars=10 µm. **(D)** VSV-G transport assay. CTRL and COPB2-KD (48% of residual protein level) cells were incubated for 1 hour at 32°C with VSV-tsO45 and cells were then shifted to 40°C for 3 hours to accumulate VSV-G (green) in the ER. The cells were then either fixed immediately (0 min) or incubated for different times (15 and 60 minutes) at 32°C in the presence of CHX and fixed. During the temperature block, both CTRL and COPB2-KD cells accumulate VSV-G into the ER. After releasing the temperature block, in both CTRL and COPB2-KD cells VSV-G reached the Golgi complex, without showing any significant delay in exiting the ER. Scale bars=10 µm. **(E)** Quantification of the ER exit of VSV-G in HeLa cells, expressed as percentage of cells with VSV-G in the ER. N=3 experiments, n=100 cells counted. Mean values ± SD. ns= not significant.

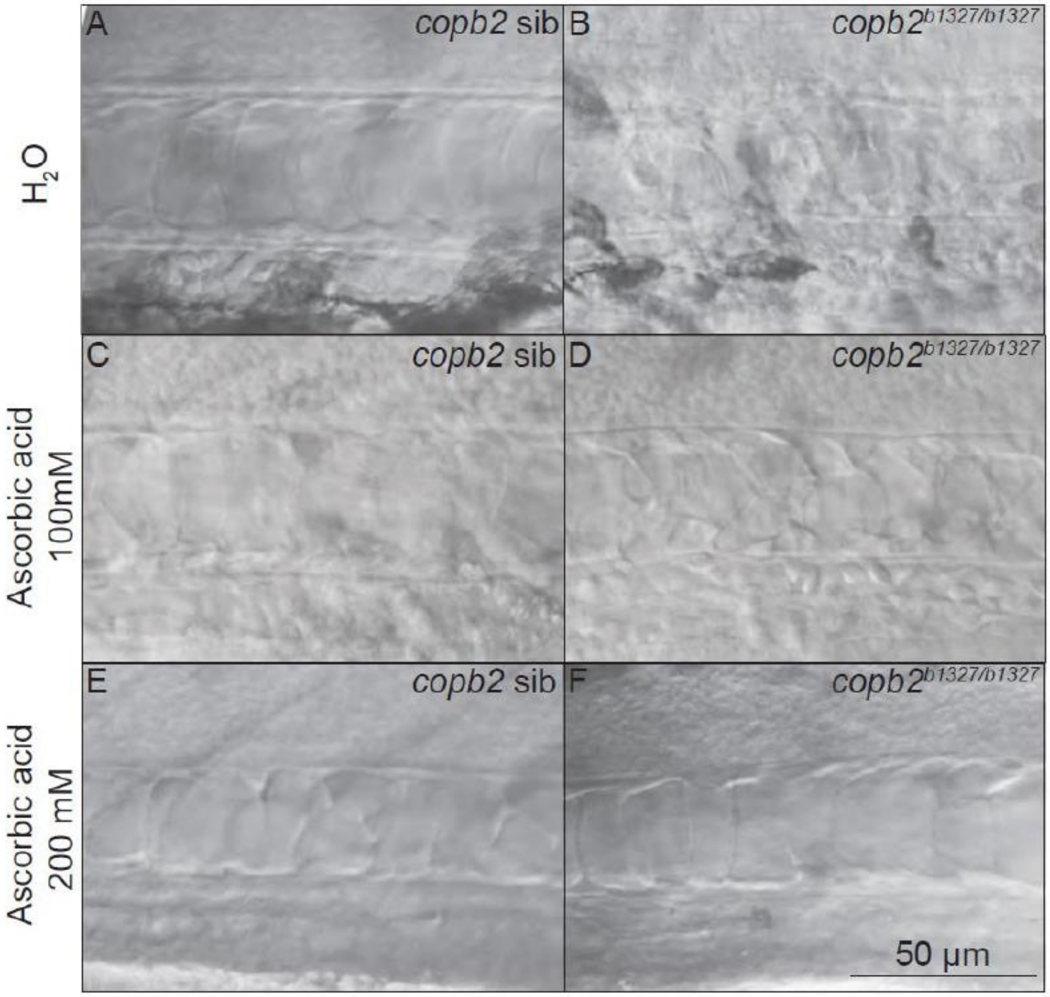


**Supplemental Figure 8. Notochord cell organization in *copb2*$^{b1327/b1327}$ mutants can be rescued by addition of ascorbic acid.** Differential interference contrast images of untreated **(A)** *copb2* siblings and **(B)** *copb2*$^{b1327/b1327}$ mutant embryos, 100 mM ascorbic acid treated **(C)** *copb2* siblings and **(D)** *copb2*$^{b1327/b1327}$ mutant embryos, and 200 mM ascorbic acid treated **(E)** *copb2* siblings and **(F)** *copb2*$^{b1327/b1327}$ mutant embryos. Anterior is to the left and dorsal to the top. Images were taken at the level of the yolk extension at 30 hpf.

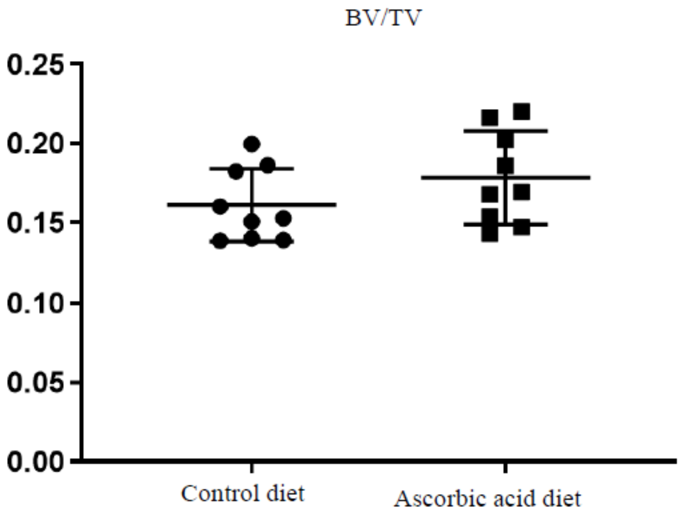


**Supplemental Figure 9. Ascorbic acid-enriched diet is associated with trend of improvement in *Copb2*$^{+/-}$ female bone mass.** In *Copb2* $^{+/-}$ female mice fed with ascorbic acid enriched diet micro CT analysis of spines showed a trend of improvement in the bone volume/total volume (BV/TV) that is not statistically significant (n=9 for each group).

| | Group | N | Mean | Std. Deviation | P value |
|---|---|---|---|---|---|
| **Total Energy (N/mm)** | WT | 7 | 12.93 | 3.79 | 0.558 |
| | Het | 9 | 13.89 | 2.62 | |
| **Plastic Displacement (mm)** | WT | 7 | 1.64 | 0.64 | 0.111 |
| | Het | 9 | 2.22 | 0.70 | |
| **Ultimate Strength (MPa)** | WT | 7 | 103.18 | 6.78 | 0.498 |
| | Het | 9 | 106.03 | 9.02 | |
| **Elastic Modulus (MPa)** | WT | 7 | 3421.04 | 538.65 | 0.955 |
| | Het | 9 | 3403.76 | 629.86 | |
| **M_Max (Adj. Ultimate Load)** | WT | 7 | 20.28 | 1.37 | 0.006 |
| | Het | 9 | 17.71 | 1.73 | |
| **Rigidity (Adj. Stiffness)** | WT | 7 | 415.95 | 50.11 | 0.018 |
| | Het | 9 | 337.42 | 63.23 | |
| **AP Diameter** | WT | 7 | 1.24 | 0.037 | 0.006 |
| | Het | 9 | 1.18 | 0.031 | |
| **CSMI** | WT | 7 | 0.12 | 0.009 | <0.0001 |
| | Het | 9 | 0.09 | 0.007 | |

**Supplemental Table 1. Summary of biomechanical testing in Copb2$^{+/-}$ mouse femurs.**

Biomechanical testing in *Copb2* $^{+/-}$ mouse femurs, showing that ultimate strength, elastic modulus and total energy were comparable to wild-type littermates, thereby supporting osteoporosis phenotype due to osteopenia, as opposed to intrinsic bone material abnormalities and increased fragility as is seen in OI mouse models.

**Members of the Undiagnosed Diseases Network**

Maria T. Acosta, Margaret Adam, David R. Adams, Pankaj B. Agrawal, Mercedes E. Alejandro, Justin Alvey, Laura Amendola, Ashley Andrews, Euan A. Ashley, Mahshid S. Azamian, Carlos A. Bacino, Guney Bademci, Eva Baker, Ashok Balasubramanyam, Dustin Baldridge, Jim Bale, Michael Bamshad, Deborah Barbouth, Pinar Bayrak-Toydemir, Anita Beck, Alan H. Beggs, Edward Behrens, Gill Bejerano, Jimmy Bennet, Beverly Berg-Rood, Jonathan A. Bernstein, Gerard T. Berry, Anna Bican, Stephanie Bivona, Elizabeth Blue, John Bohnsack, Carsten Bonnenmann, Devon Bonner, Lorenzo Botto, Brenna Boyd, Lauren C. Briere, Elly Brokamp, Gabrielle Brown, Elizabeth A. Burke, Lindsay C. Burrage, Manish J. Butte, Peter Byers, William E. Byrd, John Carey, Olveen Carrasquillo, Ta Chen Peter Chang, Sirisak Chanprasert, Hsiao-Tuan Chao, Gary D. Clark, Terra R. Coakley, Laurel A. Cobban, Joy D. Cogan, Matthew Coggins, F. Sessions Cole, Heather A. Colley, Cynthia M. Cooper, Heidi Cope, William J. Craigen, Andrew B. Crouse, Michael Cunningham, Precilla D'Souza, Hongzheng Dai, Surendra Dasari, Mariska Davids, Jyoti G. Dayal, Matthew Deardorff, Esteban C. Dell'Angelica, Shweta U. Dhar, Katrina Dipple, Daniel Doherty, Naghmeh Dorrani, Emilie D. Douine, David D. Draper, Laura Duncan, Dawn Earl, David J. Eckstein, Lisa T. Emrick, Christine M. Eng, Cecilia Esteves, Tyra Estwick, Marni Falk, Liliana Fernandez, Carlos Ferreira, Elizabeth L. Fieg, Laurie C. Findley, Paul G. Fisher, Brent L. Fogel, Irman Forghani, Laure Fresard, William A. Gahl, Ian Glass, Rena A. Godfrey, Katie Golden-Grant, Alica M. Goldman, David B. Goldstein, Alana Grajewski, Catherine A. Groden, Andrea L. Gropman, Irma Gutierrez, Sihoun Hahn, Rizwan Hamid, Neil A. Hanchard, Kelly Hassey, Nichole Hayes, Frances High, Anne Hing, Fuki M. Hisama, Ingrid A. Holm, Jason Hom, Martha Horike-Pyne, Alden Huang, Yong Huang, Rosario Isasi, Fariha Jamal, Gail P. Jarvik, Jeffrey Jarvik, Suman Jayadev, Jean M. Johnston, Lefkothea Karaviti, Emily G. Kelley, Jennifer Kennedy, Dana Kiley, Isaac S. Kohane, Jennefer N. Kohler, Deborah Krakow, Donna M. Krasnewich, Elijah Kravets, Susan Korrick, Mary Koziura, Joel B. Krier, Seema R. Lalani, Byron Lam, Christina Lam, Brendan C. Lanpher, Ian R. Lanza, C. Christopher Lau, Kimberly LeBlanc, Brendan H. Lee, Hane Lee, Roy Levitt, Richard A. Lewis, Sharyn A. Lincoln, Pengfei Liu, Xue Zhong Liu, Nicola Longo, Sandra K. Loo, Joseph Loscalzo, Richard L. Maas, Ellen F. Macnamara, Calum A. MacRae, Valerie V. Maduro, Marta M. Majcherska, Bryan Mak, May Christine V. Malicdan, Laura A. Mamounas, Teri A. Manolio, Rong Mao, Kenneth Maravilla, Thomas C. Markello, Ronit Marom, Gabor Marth, Beth A. Martin, Martin G. Martin, Julian A. Martínez-Agosto, Shruti Marwaha, Jacob McCauley, Allyn McConkie-Rosell, Colleen E. McCormack, Alexa T. McCray, Elisabeth McGee, Heather Mefford, J. Lawrence Merritt, Matthew Might, Ghayda Mirzaa, Eva Morava, Paolo M. Moretti, Marie Morimoto, John J. Mulvihill, David R. Murdock, Mariko Nakano-Okuno, Avi Nath, Stan F. Nelson, John H. Newman, Sarah K. Nicholas, Deborah Nickerson, Shirley Nieves-Rodriguez, Donna Novacic, Devin Oglesbee, James P. Orengo, Laura Pace, Stephen Pak, J. Carl Pallais, Christina GS. Palmer, Jeanette C. Papp, Neil H. Parker, John A. Phillips III, Jennifer E. Posey, Lorraine Potocki, Barbara N. Pusey, Aaron Quinlan, Wendy Raskind, Archana N. Raja, Deepak A. Rao, Genecee Renteria, Chloe M. Reuter, Lynette Rives, Amy K. Robertson, Lance H. Rodan, Jill A. Rosenfeld, Natalie Rosenwasser, Maura Ruzhnikov, Ralph Sacco, Jacinda B. Sampson, Susan L. Samson, Mario Saporta, C. Ron Scott, Judy Schaechter, Timothy Schedl, Kelly Schoch, Daryl A. Scott, Prashant Sharma, Vandana Shashi, Jimann Shin, Rebecca Signer, Catherine H. Sillari, Edwin K. Silverman, Janet S. Sinsheimer, Kathy Sisco,

Edward C. Smith, Kevin S. Smith, Emily Solem, Lilianna Solnica-Krezel, Rebecca C. Spillmann, Joan M. Stoler, Nicholas Stong, Jennifer A. Sullivan, Kathleen Sullivan, Angela Sun, Shirley Sutton, David A. Sweetser, Virginia Sybert, Holly K. Tabor, Cecelia P. Tamburro, Queenie K.-G. Tan, Mustafa Tekin, Fred Telischi, Willa Thorson, Cynthia J. Tifft, Camilo Toro, Alyssa A. Tran, Brianna M. Tucker, Tiina K. Urv, Adeline Vanderver, Matt Velinder, Dave Viskochil, Tiphanie P. Vogel, Colleen E. Wahl, Stephanie Wallace, Nicole M. Walley, Chris A. Walsh, Melissa Walker, Jennifer Wambach, Jijun Wan, Lee-kai Wang, Michael F. Wangler, Patricia A. Ward, Daniel Wegner, Mark Wener, Tara Wenger, Katherine Wesseling Perry, Monte Westerfield, Matthew T. Wheeler, Jordan Whitlock, Lynne A. Wolfe, Jeremy D. Woods, Shinya Yamamoto, John Yang, Guoyun Yu, Diane B. Zastrow, Chunli Zhao, Stephan Zuchner

### Members of the Genomics England Research Consortium

Ambrose, J. C. [1]; Arumugam, P.[1]; Baple, E. L. [1]; Bleda, M. [1]; Boardman-Pretty, F. [1;2]; Boissiere, J. M. [1]; Boustred, C. R. [1]; Brittain, H.[1]; Caulfield, M. J.[1;2]; Chan, G. C. [1]; Craig, C. E. H. [1]; Daugherty, L. C. [1]; de Burca, A. [1]; Devereau, A. [1]; Elgar, G. [1;2]; Foulger, R. E. [1]; Fowler, T. [1]; Furió-Tarí, P. [1]; Hackett, J. M. [1]; Halai, D. [1]; Hamblin, A.[1]; Henderson, S.[1;2]; Holman, J. E. [1]; Hubbard, T. J. P. [1]; Ibáñez, K.[1;2]; Jackson, R. [1]; Jones, L. J. [1;2]; Kasperaviciute, D. [1;2]; Kayikci, M. [1]; Kousathanas, A. [1]; Lahnstein, L. [1]; Lawson, K. [1]; Leigh, S. E. A. [1]; Leong, I. U. S. [1]; Lopez, F. J. [1]; Maleady-Crowe, F. [1]; Mason, J. [1]; McDonagh, E. M. [1;2]; Moutsianas, L. [1;2]; Mueller, M. [1;2]; Murugaesu, N. [1]; Need, A. C. [1;2]; Odhams, C. A. [1]; Patch, C. [1;2]; Perez-Gil, D. [1]; Pereira, M. B.[1]; Polychronopoulos, D. [1]; Pullinger, J. [1]; Rahim, T. [1]; Rendon, A. [1]; Riesgo-Ferreiro, P.[1]; Rogers, T. [1]; Ryten, M. [1]; Savage, K. [1]; Sawant, K. [1]; Scott, R. H. [1]; Siddiq, A. [1]; Sieghart, A. [1]; Smedley, D. [1;2]; Smith, K. R. [1;2]; Sosinsky, A. [1;2]; Spooner, W. [1]; Stevens, H. E. [1]; Stuckey, A. [1]; Sultana, R. [1]; Thomas, E. R. A. [1;2]; Thompson, S. R. [1]; Tregidgo, C. [1]; Tucci, A. [1;2]; Walsh, E. [1]; Watters, S. A. [1]; Welland, M. J. [1]; Williams, E. [1]; Witkowska, K. [1;2]; Wood, S. M. [1;2]; Zarowiecki, M.[1].

1. Genomics England, London, UK

2. William Harvey Research Institute, Queen Mary University of London, London, EC1M 6BQ, UK.
